# Transmembrane-domain-driven heterodimer assembly licenses integrin head-domain folding in the endoplasmic reticulum

**DOI:** 10.64898/2026.10.25.761699

**Authors:** Jakob M. Reber, Dharmendra Pandey, Hannelore Meyer, Ralph T. Böttcher, Wangqing Zhang, Peter William Krenn, Barbara Steigenberger, Dieter Langosch, Arnoud Sonnenberg, Reinhard Fässler, Shiny Shengzhen Guo

## Abstract

Integrins are a large family of α/β heterodimeric receptors whose activation is associated with the extension of the ectodomain and disruption of transient transmembrane domain (TMD) interactions. Whether the TMD interaction serves additional function(s) beyond regulating integrin activation remains unclear. Here, we report that three TMD-disruptive mutations within the integrin β1-subunit activate its function while simultaneously reducing its surface expression. While a Gly744Asn (β1-GN) and Gly744Leu (β1-GL) TMD mutation yielded viable homozygous offspring, the more disruptive Leu748Arg (β1-LR) TMD mutation caused embryonic lethality in mice and drastically reduced β1-integrin surface levels. Cell biology and biochemistry studies revealed that TMD-TMD interactions drive efficient α/β-heterodimer assembly, thereby enabling the coupled folding of the ER-luminal βI-domain. Disruption of these TMD interactions leads to endoplasmic reticulum (ER) retention and luminal domain-dependent ER-associated degradation of the monomeric, misfolded β1-subunit. Moreover, mutations that disrupt the α/β TMD interaction of multiple integrin classes in humans similarly impair integrin cell surface expression and cause severe loss-of-function diseases. Together, our findings establish that α/β TMD association serves a previously unrecognized role in integrin biogenesis by promoting βI-domain folding, preventing premature integrin activation in the ER, and ensuring the proper cell surface expression of activatable integrins.

## Introduction

Integrins mediate cell-matrix and cell-cell adhesion, which is essential for development and postnatal homeostasis. Integrins are heterodimers consisting of α- and β-subunits that associate non-covalently through their N-terminal head domains in a 1:1 complex [1]. The integrin head domain, consisting of the α-subunit β-propeller domain and the β-subunit βI-domain, harbors the ligand binding site. This head domain is followed by long, flexible leg domains, transiently interacting transmembrane domains (TMD) and short cytosolic tails (CT) that bind numerous adaptor proteins [2].

The β1-subunit pairs with 12 different α-subunits and is typically produced in excess over the α-subunits. Folding and heterodimerization of β1 occurs in the endoplasmic reticulum (ER) on a timescale of hours [3,4] and involves the engagement of chaperone proteins and folding enzymes, such as calnexin and protein disulfide isomerases [5–8]. To achieve high affinity ligand binding, integrins must undergo an activation step that leads to gross conformational changes affecting the entire molecule. An inactive integrin adopts a bent-closed conformation, in which the heterodimer is bent in the knee region of the legs and the ligand binding pocket is closed. Upon activation, the ectodomain straightens into an extended-open conformation characterized by separation of the legs, including the TMDs, and a swing out of the β-subunit hybrid domain away from the α-subunit. These rearrangements allosterically open the ligand-binding pocket to allow high-affinity ligand binding [9–11]. The importance of the ectodomain extension and TMD separation was first revealed by structural studies and later confirmed in cells using FRET-based microscopy and engineered disulfide bonds into different domains of the α- and β-subunits. These disulfide bonds were designed to impair hybrid domain swing-out, TMD separation or ectodomain extension, thereby demonstrating the essential role of the conformational changes in integrin activation [12–17].

Integrin activation is stabilized by the cytoplasmic adaptor proteins talin and kindlin, which bind to distinct sites within the β-subunit CT [18]. It is believed that the key function of the talin and kindlin association with the β-CT is to induce and/or maintain the separation of the TMD and CT, as well as to allow force transduction which stabilizes and reinforces the integrin-ligand bond [19–24]. The majority of integrins on the plasma membrane of resting cells as well as in the secretory pathway adopt the inactive, bent conformation [25]. This inactivity is mediated, at least in part, by the interaction between the long helices of the integrin α and β TMDs, which presumably contributes to the high energy barrier for transitioning between the active and inactive conformation [26–29]. The α/β-TMD interaction consists of an outer membrane clasp (OMC) and an inner membrane clasp (IMC). In the OMC, a conserved GXXXG motif in the α-TMD and its corresponding residues in the β-TMD associate through ridge-in-groove packing [30,31]. The IMC is mediated by hydrophobic stacking of the α/β-TMD and a salt bridge formed between the conserved arginine residue of the α-CT (GFFK<u>R</u>) and aspartic acid (D) residue in the β-CT [30]. Mutations of critical residues in either the OMC or the salt bridge of integrin β3 weaken α/β TMD interactions and result in elevated fractions of active integrins on cells *in vitro* [30–33].

To investigate the functional significance of the regulation of TMD-TMD interaction *in vivo*, we generated knock-in mice carrying point mutations in the OMC of the β1-TMD. Specifically, two mutations (β1-G744N, β1-G744L, β1-GN and β1-GL, respectively) were introduced to alter the packing environment within the β1-TMD. Additionally, we substituted the hydrophobic leucine at position 748 in the β1-TMD with a positively charged arginine residue, which is predicted to disrupt helix-to-helix alignment by shortening the β-TMD (β1-L748R; β1-LR) and to destabilize the clasp with the α-TMD helix. Whereas the β1-GN and β1-GL mouse strains produced viable homozygous offspring, the β1-LR mouse strain arrested development shortly after implantation. Biochemical and cellular assays revealed that the β1-LR substitution disrupts the α/β TMD association, which is critical for the proper assembly and folding of the α-subunit β-propeller domain and the β-subunit βI-domain.

## Results

### The integrin TMD clasp is required for integrin surface expression

The dissociation and association of the integrin α/β TMDs have been shown to regulate the activity state of the β2- and β3-class integrins [27, 30, 32]. To examine whether this mechanism is conserved in members of the large β1-class integrins, we introduced mutations at corresponding positions in the β1-TMD that are known to disrupt the α/β TMDs in β3 integrins (G708N, G708L and L712R, Figure S1A [32,34,35]. We first assessed the effects of these mutations in the β1 subunit (β1-G744N, β1-G744L and β1-L748R, Figure S1A) on heterodimerization with the α5 subunit TMD using the split β-lactamase-TM (BLaTM) assay [36]. In this assay, antibiotic resistance is conferred to *Escherichia coli* when the α5-TMD and β1-TMD fused to the N- and C-terminal β-lactamase (N-BLa, C-BLa), interact to reconstitute β-lactamase activity. Regardless of the fusion orientation, all three β1-TMD mutations strongly decreased ampicillin resistance of bacteria (Figure 1A and Figure S1B-D) indicating impaired heterodimerization with the wild type α5-TMD.

**FIGURE 1.**
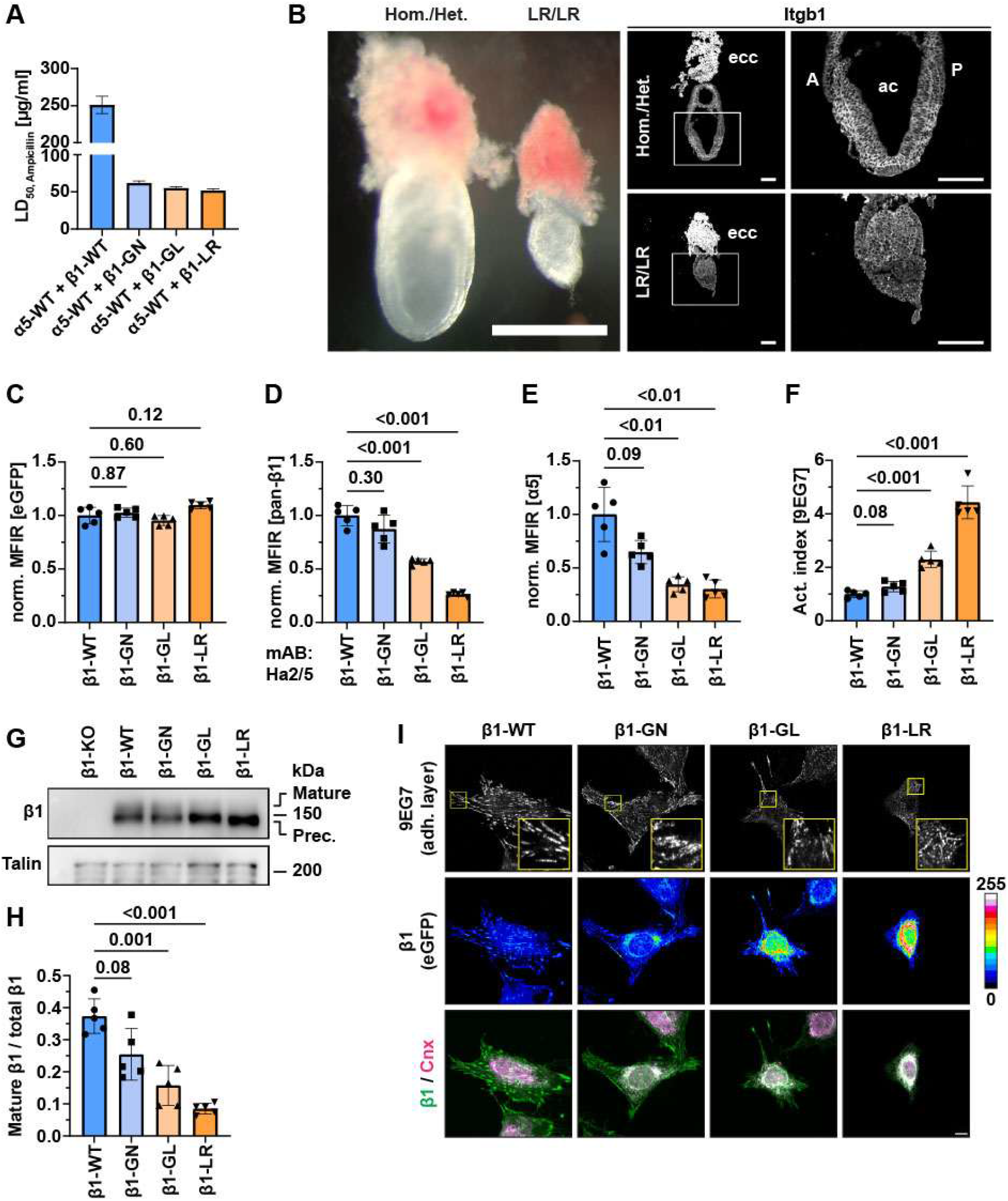
Functional consequences of β1 TMD domain mutations. (A) TMD-TMD interactions between wild-type or mutant β1 subunits with the α5 subunit measured using the β-lactamase (Bla) TM assay. Heterotypic interactions were assessed by expressing the TMDs of β1 and α5 subunits fused to the N-(N-Bla-β1) and C-terminal (C-Bla-α5) β-lactamase fragments, respectively, in *Escherichia coli*. Antibiotic resistance as a measure of TMD-TMD affinity is indicated as LD50 values obtained from global fits of pooled OD600 measurements from an ampicillin titration series (n=6 biological replicates). (B) Whole-mount widefield and confocal microscopic images of E7.5 wild-type and β1-LR embryos stained for β1 (n=3). Note the defective germ layer formation, undefined A-P axis and failed cavitation in the β1-LR embryos. ecc: ectoplacental cone; ac: amniotic cavity; A: anterior; P: posterior; Hom./het.: Putative homozygous/heterozygous embryo. (C-F) Flow cytometry of β1-KO cells stably expressing the indicated GFP-tagged β1-subunits (n=5). Activation index was calculated as the ratio of 9EG7 signals to total β1 signals (antibody clone: Ha2/5). MFIR: mean fluorescence intensity ratio. (G,H) Representative Western blot of lysates from cells expressing the indicated β1 subunits (G) and densitometric quantifications of mature (125 kDa) versus total β1 integrin levels from 4 independent experiments (H). Talin served as loading control. (I) Representative immunofluorescence images of FN-seeded cells expressing the indicated GFP-tagged β1-subunits and immunostained for the β1 integrin activation-reporting antibody 9EG7 and the ER marker calnexin (Cnx). 9EG7 signals were collected from confocal slices at the adhesive substrate. Boxes highlight magnified 9EG7-positive FAs. Due to differences in spreading and cell height, contrast in overlay is enhanced for maximal visibility and not quantitative. Scale bar, 10 µm. Statistical analysis was performed using Brown-Forsythe and Welch ANOVA with Dunnett T3’s post-hoc test. Data are presented as mean±SD.

Next, we introduced the G744N, G744L and L748R mutations (β1-GN/-GL/-LR, respectively) into the *Itgb1* gene of mice using embryonic stem (ES) cells [37]. Mice heterozygous or homozygous for the β1-GN and β1-GL substitutions were born at normal Mendelian ratios (Figure S1E,F), displayed no obvious abnormalities and had a normal life span. While mice heterozygous for the β1-L748R are also born at normal ratios and did not display apparent abnormalities, intercrosses between β1-LR heterozygotes failed to produce live homozygous offspring (Figure S1G), indicating that the lethal phenotype is recessive. Timed mating combined with PCR genotyping revealed that at E7.5 homozygous β1-LR embryos were present at expected Mendelian ratios (Figure S1H). These embryos consisted of either ectoplacental cones without embryo proper or with small, malformed embryonic structures that lacked an amniotic cavity, clear germ layers or the anterior-posterior body axis (Figure 1B).

To further investigate the cellular consequences of the β1-TMD mutations, we retrovirally transduced β1-integrin-null (β1-KO) mouse fibroblasts with cDNAs encoding eGFP-tagged human β1-WT [38], as well as the -GN, -GL and -LR mutants. Cells with comparable eGFP expression levels were isolated by sorting (Figure 1C). Despite similar eGFP levels, total β1 surface expression remained unchanged in fibroblasts expressing β1-GN, reduced in β1-GL and barely detectable in β1-LR. Concomitant with this progressive decrease in β1 surface levels, the levels of the α5- and the α6-subunits also gradually declined (Figure 1D,E and Figure S1I). Western blotting revealed a corresponding decrease in mature β1-integrin (representing cell surface β1-integrin) in lysates from β1-GN, β1-GL and β1-LR cells, accompanied by a relative increase in the β1-integrin precursor (representing the ER-resident β1-integrin; Figure 1G,H). Treatment of lysates from cells expressing equal amounts of β1-WT or β1-LR (Figure S1J) with Endo H or PNGase F confirmed the distinct glycosylation status of the β1 precursor and mature forms. The ER-localized β1 precursor carried high-mannose glycans and was sensitive to both Endo H and PNGase F, whereas the mature β1 subunit at the cell surface, in which high-mannose glycans had been processed into complex-type glycans in the Golgi, was sensitive only to PNGase F (Figure S1K). Finally, immunostaining for β1 showed a loss of β1-integrin from focal adhesions (FAs), impaired spreading on fibronectin (FN) and increased colocalization of β1 with the calnexin-positive (Cnx^+^) ER in β1-GN, -GL and -LR cells compared with β1-WT (Figure 1I).

To assess the activation state of mutant β1-integrins at the cell surface, we determined their activation index, defined by the ratio of active to total β1-integrin levels. Active β1-integrins were measured using the monoclonal antibodies (mAbs) 9EG7 (reporting ectodomain extension), Huts21 (reporting hybrid domain swing-out) and the 12G10 (reporting headpiece opening). All β1-TM mutations exhibited elevated activation indexes, with β1-LR showing the largest increase (Figure 1F and Figure S1L,M). These findings indicate that the α/β TM clasp both promotes ER exit and curbs integrin activation on the cell surface.

### The β1-integrin TMD mutants fail to heterodimerize with α-integrin subunits

The ER retention and reduced surface expression of β1-GN, β1-GL and β1-LR integrins could potentially result from improper insertion of the mutant TMDs into the ER membrane. To test this possibility, we performed membrane fractionation experiments, which revealed that β1-WT as well as β1-GN, -GL and -LR integrins were almost exclusively present in the pellet/membrane fraction from both high salt and carbonate extractions (Figure 2A and Fig S2A), suggesting that the TMD mutations do not impair membrane insertion.

**FIGURE 2.**
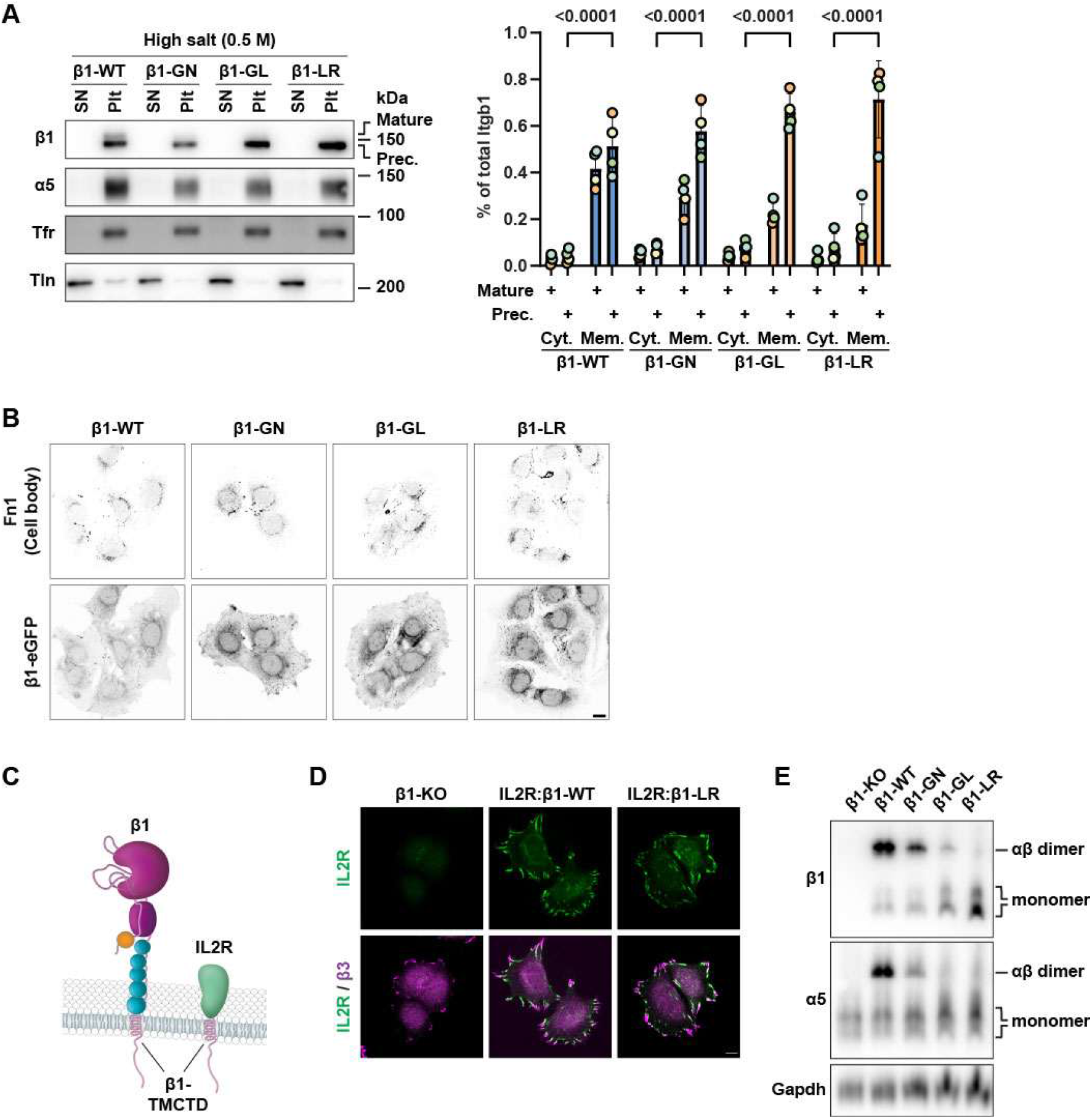
Mutations in the β1 TMD impair the heterodimerization with α-subunits. (A) Western blots of high-salt-extracted membrane and cytoplasmic fractions from cells expressing the indicated β1-subunits, stained for β1 and α5. transferrin receptor (Tfr) and talin (Tln) served as membrane and cytoplasmic controls, respectively (n=4). SN: supernatant; Plt: pellet. Densitometric quantification of precursor and mature β1-integrin in the cytoplasmic and total membrane fraction is shown on the right as a percentage of total β1 integrin from n=4 independent experiments. (B) Representative immunofluorescence images of serum-seeded cells expressing the indicated β1-subunits and stained for FN. (C) Schematic of the β1-subunit and the chimeric IL2R:β1, in which the IL2R ectodomain is fused to the β1-WT or β1-LR TMD and CTD of β1. (D) Representative immunofluorescence images of FN-seeded β1-KO cells and β1-KO cells reconstituted with IL2R:β1-WT or IL2R:β1-LR, stained for the IL2R ectodomain (green) and the β3-subunit (magenta) (n=3). (E) BN-PAGE followed by Western blotting of lysates from cells expressing indicated β1-subunit, probed with antibodies against β1 and α5. Gapdh served as loading control, n=3.

Next, we tested whether premature activation of the β1-GN, β1-GL and β1-LR integrins promotes FN binding in the ER and thereby prevents the packaging of potentially oversized integrin-FN complexes into transport vesicles. Immunostaining for FN showed perinuclear localization of endogenous FN in freshly seeded cells. Overall, cells expressing β1-GN, -GL and -LR exhibited perinuclear localization of FN comparable to that in β1-WT cells (Figure 2B), indicating that FN is not co-retained with mutant β1 integrins in the ER.

To investigate whether the low surface expression of the mutant β1-TMDs was due to a trafficking defect through the secretory pathway, we constructed chimeric proteins by fusing the cytosolic tail and TMD of either β1-WT or β1-LR subunits to the extracellular domain of the interleukin-2 receptor α (IL2R, Figure 2C), and expressed these chimeric proteins in β1-KO fibroblasts. Flow cytometry measurements revealed that both IL2R:β1-WT and IL2R:β1-LR chimeras were expressed at similar levels on the cell surface but did not restore the surface expression of the integrin α-subunits (Figure S2B,C). Immunostainings further demonstrated that both integrin-chimeric constructs localized to the rear of β3 integrin-positive FAs (Figure 2D), indicating that the LR mutation alone is insufficient to drive ER retention of β1-LR. To determine whether retention of β1-LR, as well as that of β1-GN and β1-GL, results from disrupted pairing with α-subunits, we performed blue native polyacrylamide electrophoresis (BN-PAGE) followed by Western blotting. β1-WT predominantly formed heterodimers, whereas heterodimerization of β1-GN, β1-GL and β1-LR integrins with the α5-subunit was impaired to varying degrees, with the most severe defect observed for β1-LR (Figure 2E). Proteomic analysis of immunoprecipitates further confirmed reduced β1-L748R heterodimerization with the α5 subunit, and additionally revealed diminished interactions with the α3, α6 and α7 subunits (Figure S2D). Together, these findings indicate that activating mutations in the β1-integrin TMD impair heterodimerization of integrins, which is required for efficient surface expression.

### The β1-subunit I-domain of β1-L748R induces ER retention

The leucine residue at position 979 (L979) in the α5-subunit TMD corresponds to αIIb-M987, which has been reported to interact with the positively charged arginine residue in β3-L712R and, based on sequence homology and AlphaFold prediction, the corresponding residue in β1 is L748R (Figure S1A and Figure S2E) [30, 39]. To test whether substituting L979 with a negatively charged aspartic acid (α5-LD, Figure 3A) could promote pairing with β1-LR and rescue its surface expression, we co-expressed the α5-LD and β1-LR subunits in fibroblasts lacking both α5 and β1 subunits (DKO cells). Surface expression and ER retention were compared to fibroblasts expressing β1-WT with either α5-WT or α5-LD, and to cells expressing β1-LR with α5-WT. After sorting for integrin-positive populations, all cell lines showed comparable mRNA expression levels (Figure S3A).

**FIGURE 3.**
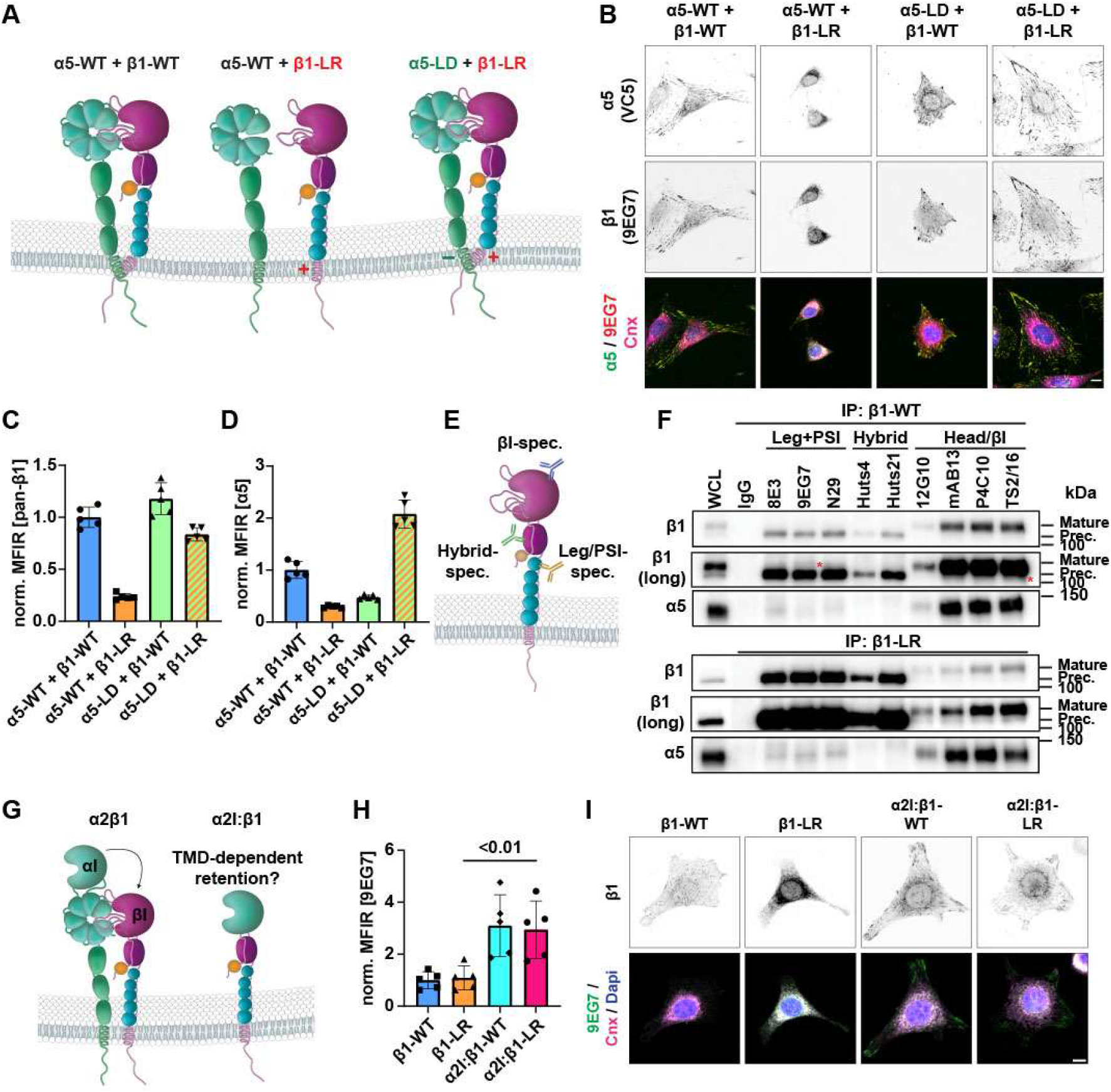
The βI-domain of β1-LR causes the ER retention. (A) Schematic of α5β1 integrins with or without mutations in the β1 subunit (β1-LR) or α5-subunit (α5-LD). (B) The indicated α5- and β1-subunits were expressed in FN-seeded α5/β1 double knockout (DKO) cells and immunostained for α5 (VC5), β1 (9EG7) and the ER marker calnexin (Cnx). Scale bar: 10 µm. (C,D) Flow cytometry-based measurements of MFIR of α5 (IIA1) and β1 (Ha2/5) surface expression in DKO cells reconstituted with the indicated α5- and β1-subunits (n=5). MFIR values are normalized to cells expressing α5-and β1-WT-subunits. Statistical analysis was performed using repeated measures one-way ANOVA with Tukey’s post-hoc test. Data are presented as mean±SD. (E) Schematic of antibody binding sites on the β1-subunit. (F) Western blot of anti-β1 immunoprecipitates from lysates of cells expressing human β1-WT or β1-LR. Immunoprecipitations were performed with control IgG and the indicated mAbs (n=3). Weak bands, potentially corresponding to small amounts of either the mature or precursor forms of β1 integrin, visible only upon prolonged exposure, are indicated by red asterisks. (G) Schematic of a βI-domain-swap with the αI domain of the α2-subunit (α2I). (H) MFIR of the 9EG7 epitope measured by flow cytometry on cells expressing β1-WT, β1-LR, α2I:β1-WT, or α2I:β1-LR (n=5), normalized to β1-WT expressing cells. Statistical analysis was performed using repeated measures ANOVA with Dunnett’s post-hoc test. Data are presented as mean±SD. (I) Representative immunofluorescence images of FN-seeded cells expressing β1-WT, β1-LR, α2I:β1-WT, or α2I:β1-LR, immunostained for β1 (4B7R: black and white; 9EG7: overlay) and calnexin (Cnx), and counterstained with DAPI. Scale bar: 10 µm.

Upon seeding on FN, DKO fibroblasts expressing the untagged β1-LR with α5-WT failed to spread, assembled few and small FAs and retained most of the α5-WT and the β1-LR subunits in the Cnx^+^ ER, with the latter exposing the 9EG7 activation epitope (Figure 3B). Flow cytometry analyses confirmed reduced surface levels of α5 and β1 integrin, as well as other α-integrin subunits such as α6 that heterodimerize with β1 (Figure 3C,D and Figure S3B). Fibroblasts expressing β1-WT with α5-LD spread less on FN, formed few FAs containing α5-LD/β1-WT integrins, and retained some α5-LD in the perinuclear region (Figure 3B) explaining the diminished surface expression (Figure 3D). In contrast, the β1-WT subunit was not retained in the ER in these cells (Figure 3C) and reached the cell surface upon heterodimerization with endogenous α-subunits such as α6, whose surface expression increased in the absence of the α5-WT subunit (Figure S3B).

In sharp contrast, co-expression of the α5-LD and β1-LR restored cell spreading to levels comparable to those of α5-WT/β1-WT cells and produced comparably high numbers of FAs and cell-centered, elongated fibrillar adhesions (Figure 3B). Because only the α5-LD subunit can efficiently interact with the β1-LR subunit, surface levels of the α5-LD were high, while endogenous α6 levels on the cell surface were reduced in α5-LD/β1-LR cells (Figure 3D and Figure S3B). Assessment of the integrin activation index using 9EG7 for β1 and SNAKA51 for α5 showed reduced activation of the α5-LD/β1-LR integrin (Figure S3C,D), consistent with stabilization of the heterodimer by a TMD ionic bond that restrains activation. These results demonstrate that artificial clasping of the TMD of the α5- and β1-subunit facilitates integrin heterodimer formation and promotes integrin surface expression.

To identify the misfolded/unfolded domain responsible for ER-retention of β1, we expressed equal amounts of full length human β1-WT or β1-LR integrin in β1-KO fibroblasts. The integrins were subsequently immuno-precipitated from cell lysates using well-characterized, conformation-specific β1 mAbs, and epitope-availability was assessed by Western blotting with a pan-human-β1 integrin antibody (Figure 3E). In β1-WT-expressing fibroblasts, leg- and hybrid-domain-specific mAbs primarily precipitated precursor β1 along with minor amounts of the α5 subunit, likely representing its ER-associated precursor form. In contrast, βI-domain-specific mAbs precipitated the mature form of β1, which co-precipitated with substantial levels of α5 (Figure 3F). In β1-LR expressing fibroblasts, leg- and hybrid-domain-specific mAbs precipitated large amounts of β1 precursor, whereas βI-domain-specific antibodies precipitated only small amounts of the mature β1-subunits (Figure 3F and Figure S3E). These findings suggest that ER-retention of β1-LR is associated with a misfolded βI-domain.

To investigate this possibility, we exchanged the βI-domain in both β1-WT and β1-LR with the structurally homologous, spontaneously folding αI-domain of the α2-integrin subunit [40], generating the chimeric constructs α2Ι:β1-WT and α2Ι:β1-LR (Figure 3G). These chimeric β1 subunits were then expressed in β1-KO fibroblasts. Flow cytometry analysis using the 9EG7 antibody revealed comparable surface levels of α2Ι:β1-WT and α2Ι:β1-LR, both of which were substantially higher than those of β1-LR, indicating that substitution with the α2I-domain restores surface expression despite the TMD-L748R mutation (Figure 3H and Figure S3F). The α5-subunit was not detectable at the surface of cells expressing α2I:β1-WT or α2I:β1-LR (Figure S3G). Both surface-expressed α2Ι:β1-WT and α2Ι:β1-LR localized to FAs and showed only minimal retention in the ER (Figure 3I). Notably, however, their surface levels were significantly lower than those of β1-WT, likely owing to rapid internalization followed by lysosomal degradation (Figure S3H,I). Together, these findings further implicate the βI-domain as a key determinant of β1-LR ER retention.

Finally, we examined whether the βI-domain folding and ER export of β1-LR depend on the interaction with the α-subunit head domain. To this end, we fused the headpiece of the α5-subunit, comprising the β-propeller and thigh domains, to the N-terminus of β1-WT or β1-LR subunit (Figure S3J). The fusions were generated either with a non-cleavable serine-glycine (SG)-containing linker peptide (α5-SG-β1-WT; α5-SG-β1-LR), or with a self-cleaving 2A peptide (α5-2A-β1-WT; α5-2A-β1-LR), which allows dissociation of the α-subunit head domain. Importantly, 2A peptide-mediated cleavage of the α-subunit head domain from the α5-2A-β1-WT polypeptide releases a free β1-WT subunit that can form heterodimers with endogenous α-subunits in the ER and subsequently reach the cell surface. Therefore, to distinguish the effects of headpiece association from those of β1 heterodimerization, the α5-2A-fusion proteins were expressed in DKO cells and their ability to form α5β1 heterodimers assessed by FACS using the β-propeller domain-specific IIA1 mAb. Cells expressing the α5-headpiece cleaved from β1-WT or β1-LR displayed comparably low surface levels of the IIA1 epitope. In contrast, cells expressing the α5-headpiece domain directly linked to β1-WT or β1-LR (α5-SG-β1-WT; α5-SG-β1-LR) exhibited comparably high surface levels of IIA1 (Figure S3K). These findings indicate that physical association of the α5-headpiece with the βI-domain promotes βI-domain folding of β1-LR, thereby facilitating its ER export to the cell surface.

### The β1-LR I-domain displays increased free cysteines

The data so far suggest that the folding state of the βI-domain causes retention of β1-LR integrins in the ER. Because protein folding in the ER often involves the formation of disulfide bonds, we compared the oxidation status of cysteines in the βI-domain of β1-WT and β1-LR, using a stable isotope differential alkylating approach. This method involves stepwise labeling of reduced and oxidized cysteines with light (d_o_) and heavy (d_5_) deuterium-labeled *N*-ethylmaleimide (NEM), allowing comparative quantification of the oxidation state of the cysteine residues between the β1-WT and β1-LR subunits. Our analysis quantified 40 out of 56 cysteines in the β1-subunit. Among these, cysteine-207 (C207) and −301 (C301) showed significantly higher levels of their free (reduced) forms in β1-LR than in β1-WT, which also exhibited high levels of free C207 and C301 likely owing to its overproduction [4] (Figure 4A). Notably, both C207 and C301 reside within the βI-domain. In the mature β1-subunit C207 forms a disulfide bond with C213 and contributes to the specificity determining loop (SDL), which is essential for ligand binding but not required for the assembly of α5β1 [41]. C301, which pairs with C261 in the mature β1 subunit, has not been functionally investigated so far (Figure 4B) [42]. However, the importance of the C261-C301 disulfide bond for βI-domain stability is underscored by sequence alignments, showing its conservation across all β-subunit paralogues in all species examined (Figure S4A-C). To assess whether the increased levels of free C207 and C301 in β1-LR are associated with increased chaperones binding, human β1-LR, mouse β1-LR and their respective β1-WT counterparts were expressed in β1-KO fibroblasts, precipitated with specific antibodies, and analyzed by MS for coprecipitated proteins. Protein disulfide isomerases P4hb and Pdia3 (Figure S5A) as well as Pdia6, Calnexin (Canx), Hspa5 and the UDP-glucose glycoprotein glucosyltransferase Uggt1 were enriched in β1-LR compared with β1-WT (Figure S5B).

**FIGURE 4.**
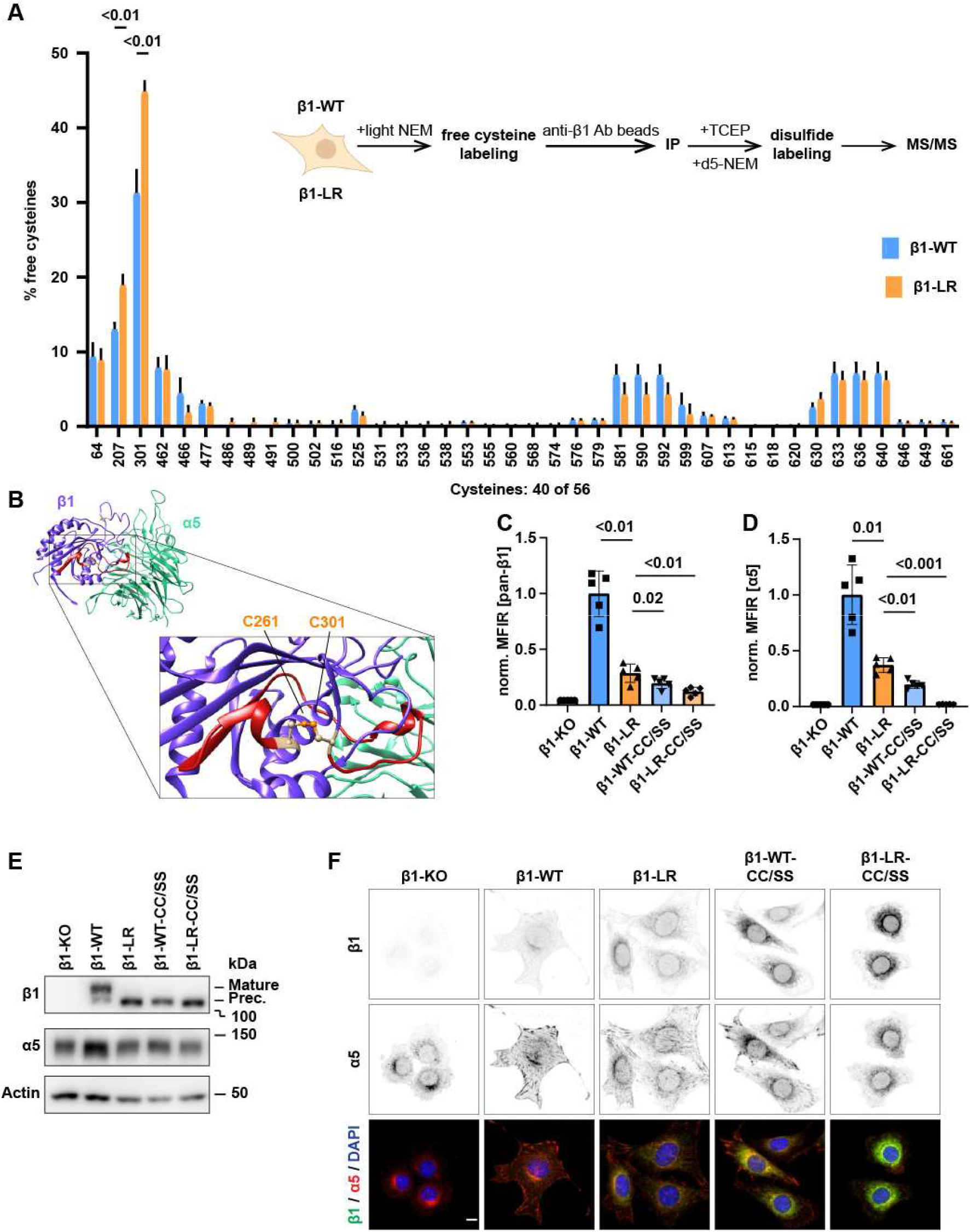
The oxidation state of cysteines in the βI domain is required for α/β heterodimerization and ER release. (A) Average of MS1 and MS2 ratio between reduced and total (reduced + oxidized) cysteine residues in β1-WT and β1-LR (n=4 biological replicates, with n=3 technical replicates for each biological replicate). Statistical analysis was performed using multiple t-tests with Dunnett T3’s post-hoc test. Data are shown as mean±SD. (B) Ribbon diagram of the α5β1 head domain (PDB: 3vi4). The position of C261 and C301 are highlighted and the loop nestling in a cavity of the α5 propeller domain is shown in red. (C,D) Flow cytometry-based measurements of β1 and α5 surface expression on KO cells expressing human β1-WT or β1-LR without or with substitution of C261 and C301 for serine (S). MFIR is normalized to β1-WT cells. Statistical analysis was performed using repeated measures one-way ANOVA with Dunnett’s post-hoc test. Data are shown as mean±SD. (E) Representative Western blot of cell lysates derived from cells expressing β1-WT or β1-LR without or with substitution of C261 and C301 to CC/SS. (F) Representative immunofluorescence images of FN-seeded β1-KO cells and β1-KO cells reconstituted with indicated β1 variants stained for β1 and α5. Scale bar is 10 µm.

To evaluate the importance of the C261-C301 disulfide bond in β1-integrin folding and surface expression, we generated expression constructs carrying the C261S and C301S substitutions in β1-WT and β1-LR (referred to as CC/SS) and introduced them into β1-KO cells via retroviral transduction. In cells expressing β1-WT-CC/SS, surface levels of β1 and α5 were lower than in cells expressing β1-LR, with an even greater reduction observed in cells expressing β1-LR-CC/SS (Figure 4C,D). Western blot analysis showed that the mature β1-subunit was barely detectable in cell lysates expressing β1-LR, β1-WT-CC/SS and β1-LR-CC/SS, whereas levels of the precursor were elevated compared to β1-WT cells (Figure 4E). Immunostaining of FN-seeded cells revealed strong β1-integrin signals in FAs and weak ER staining in β1-WT expressing cells. In contrast, β1-integrin staining in FAs was reduced and progressively increased in the ER of cells expressing β1-LR, β1-WT-CC/SS, and β1-LR-CC/SS (Figure 4F). Together, these data indicate that formation of the C261-C301 disulfide bond within the βI-domain of β1 integrin is essential for βI-domain stability and surface expression of β1 integrins.

### β1-LR and unpaired β1-WT are ERAD-L substrates

The ER-associated protein degradation (ERAD) pathway recognizes and retro-translocates terminally misfolded proteins from the ER into the cytoplasm for proteasomal degradation [43]. To determine whether the misfolded β1-LR is targeted for proteasomal or lysosomal degradation, we performed pulse-chase experiments. β1-KO fibroblasts expressing either β1-WT or β1-LR were labeled with [^35^S]methionine/cysteine for 30 minutes to generate a pulse of newly synthesized proteins. Cells were then treated with the proteasome inhibitor MG132 or the lysosome inhibitor Bafilomycin A1 (BafA1), chased with excess unlabeled methionine/cysteine, lysed, and immunoprecipitated using an anti-β1-integrin polyclonal antiserum. The stability of β1 subunits over time was then assessed by SDS-PAGE and autoradiography.

In mock-treated control samples, most of the β1-WT precursor was converted to the mature form during a 7-hour chase period, whereas β1-LR subunit remained predominantly in its precursor form (Figure 5A,B). BafA1 treatment had no apparent effect on mature β1-WT levels, likely due to high stability of integrins at the cell surface [44]. In β1-LR expressing cells, however, BafA1 treatment resulted in the accumulation of several less prominent, slower migrating β1-LR species, one of which may correspond to the mature β1 form. By contrast, MG132 treatment led to a marked accumulation of the β1-LR precursor and the appearance of the β1-WT precursor. Together, these findings strongly suggest that misfolded β1-LR and excess unpaired β1-WT subunits are retained in the ER and subsequently targeted for proteasomal degradation.

**FIGURE 5.**
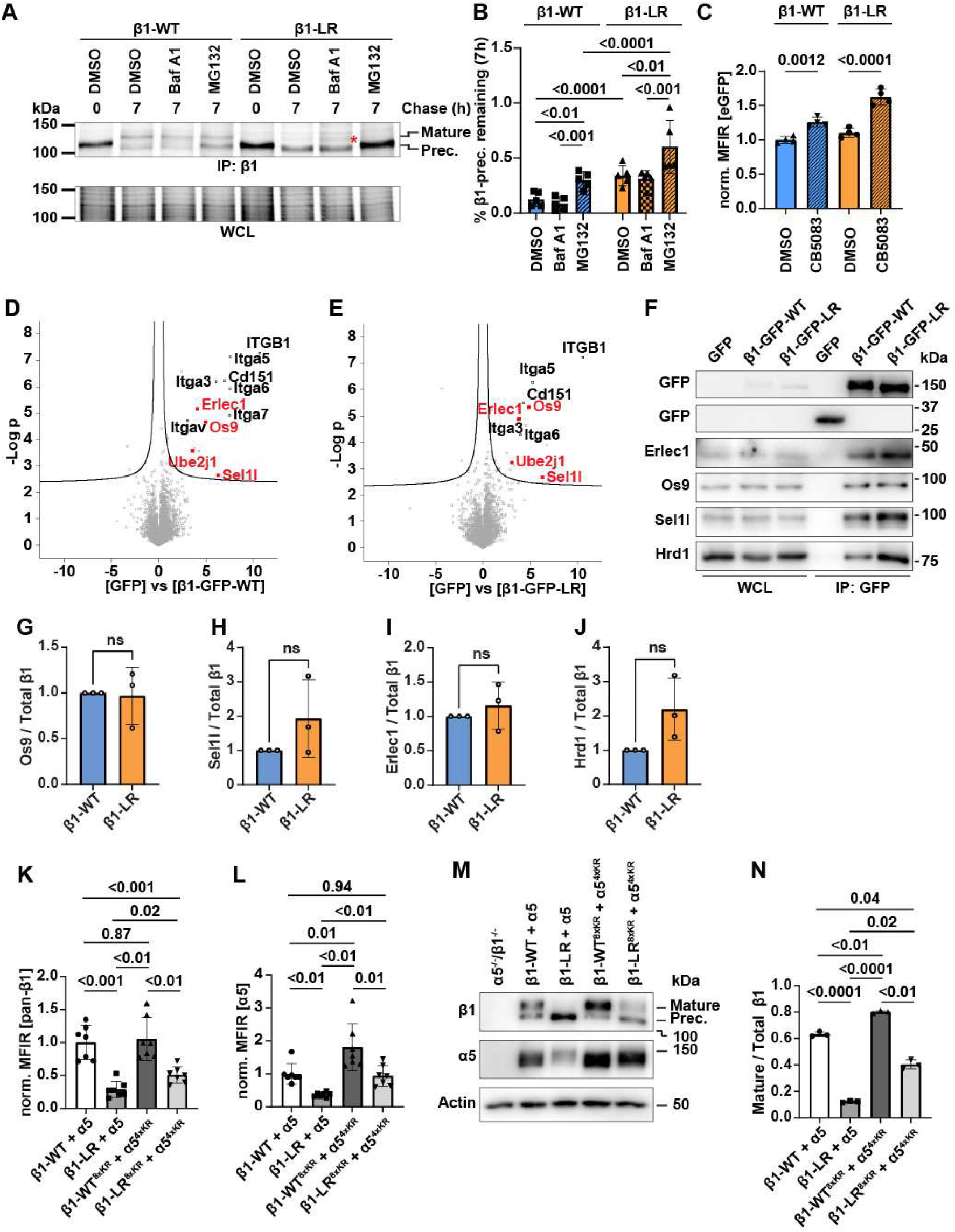
Monomeric β1-WT and β1-LR subunits are substrates of ERAD-L. (A) Representative pulse-chase analyses of β1-integrin maturation in cells expressing β1-WT or β1-LR integrins. Cells were metabolically labelled with [^35^S]-methionine/cysteine and chased for the indicated time points in the absence or presence of proteasome (MG132) or lysosome inhibitor (BafA1), followed by immunoprecipitation. (B) Densitometric quantification of precursor β1-integrin as a percentage of total β1 integrin from the pulse-chase autoradiograms (n=5). Statistical analysis was performed using two-way ANOVA with Tukey’s post-hoc test. Data are presented as mean±SD. (C) Flow cytometry-based measurements of MFIR of cells expressing the indicated GFP-tagged β1-subunits treated with DMSO or p97/Vcp inhibitor CB5083. MFIR values are normalized to DMSO-treated β1-WT cells. Statistical analysis was performed using repeated measures one-way ANOVA with Bonferroni’s post-hoc test. Data are presented as mean±SD. (D,E) Volcano plot of proteins immunoprecipitated from CB5083-treated β1-KO cells reconstituted with β1-GFP-LR or GFP-only (n=4 biological replicates). Statistical analysis was performed using two-sided permuted t-test (250 randomizations). (F) Western blot of anti-GFP immunoprecipitates and whole cell lysates (WCL) from CB5083-treated β1-KO cells reconstituted with GFP-only or GFP-tagged β1-WT, or β1-LR. (G-J) Densitometric quantifications of co-precipitated Os9, Sel1l, Erlec1 and Hrd1 as a fraction of total precipitated β1 (n=3 independent experiments). (K,L) Flow cytometry-based MFIR of total β1 (Ha2/5 antibody) and total α5 (IIA1 antibody) in DKO cells retrovirally transduced with β1-WT^WT/8xKR^-IRES2-α5^WT/4xKR^ and β1-LR^WT/8xKR^-IRES2-α5^WT/4xKR^, respectively. MFIR is normalized to β1-WT cells. Statistical analysis was performed using repeated measures one-way ANOVA with Tukey’s post-hoc test. Data are shown as mean±SD. (M) Representative Western blot of WCL from DKO cells transduced with β1-WT^WT/8xKR^-IRES2-α5^WT/4xKR^ and β1-LR^WT/8xKR^-IRES2-α5^WT/4xKR^. (N) Densitometric measurement of mature β1-integrin from Western blots shown in (M) as a percentage of total β1 integrin (n=3). Statistical analysis was performed using repeated measures one-way ANOVA with Tukey’s post-hoc test. Data are shown as mean±SD.

The translocation of terminally misfolded β1-LR and excess unpaired β1-WT from the ER to the proteasome suggests that both are targeted for degradation via ERAD, in which substrates are retro-translocated from the ER membrane, polyubiquitinated by the Sel1l-Hrd1 E3 ubiquitin ligase complex, extracted into the cytoplasm by the AAA+-ATPase valosin-containing protein (Vcp/p97) and degraded by the proteasome [45–48]. Consistently, chemical inhibition of Vcp/p97 with CB-5083 increased total protein levels in β1-KO fibroblasts expressing GFP-tagged β1-WT as well as β1-LR (Figure 5C), indicating that ERAD mediates the clearance of misfolded β1-LR (quality control) and excess unpaired β1-WT subunits (quantity control) [49].

If ERAD is indeed involved in clearing β1-LR and unpaired β1-WT with misfolded βI-domains, ERAD-L components should interact with the β1-subunits. To test this hypothesis, we expressed GFP-tagged β1-WT, β1-LR or GFP alone in β1-KO cells, chemically inhibited Vcp/p97 with CB-5083 to enrich for misfolded ERAD-target proteins in the ER, performed GFP immunoprecipitation and analyzed the co-immunoprecipitate by MS. The experiment revealed that comparable amounts of Os9, Erlec1 and Sel1l co-precipitated with both β1-WT and β1-LR (Figure 5D,E). Western blot analysis confirmed the MS results and additionally showed that both β1-WT and β1-LR associate with Hrd1 (Figure 5F-J). These findings demonstrate that β1-LR is targeted for ERAD-L-mediated degradation and identify excess β1-WT as a novel endogenous substrate of ERAD.

A key function of ERAD is to polyubiquitinate misfolded substrates through the Hrd1/Sel1l complex. Hence, expression of non-ubiquitinable integrin subunits should impair ERAD and increase β1-LR expression. To test this hypothesis, we substituted four lysine residues in the CT of the α5-subunit and eight in the CT of the β1-subunit with arginine residues creating β1-WT^8xKR^, β1-LR^8xKR^ and α5^4xKR^ mutants. These mutants were expressed in various combinations in α5^-/-^β1^-/-^ DKO fibroblasts. Flow cytometry analysis showed that cells expressing α5^4xKR^/β1-WT^8xKR^ subunits, as well as cells expressing the α5/β1-WT subunits had similar levels of β1 integrins on their surface. In contrast, expression of α5^4xKR^/β1-LR^8xKR^ led to approximately a two-fold increase in β1 integrin surface levels compared to cells expressing the α5/β1-LR integrin (Figure 5K,L). Western blots analysis confirmed that the level of the mature β1 subunit was increased in α5^4xKR^/β1-LR^8xKR^ expressing cells relative to α5/β1-LR, while the precursor form decreased, resulting in a quantitative shift in the ratio of mature to total β1 integrins (Figure 5M,N).

### Disruptive disease mutations in TMDs reduce integrin surface expression

Several missense mutations in the TMDs of the integrin α- and β-subunits (Figure S1A) have been linked to inherited diseases that impair integrin activity and/or surface expression [50–57]. To investigate whether these mutations, like the β1-LR substitution, disrupt α/β TMD interaction and reduce integrin surface levels, we analyzed representative disease-associated TMD mutations across different integrin subfamilies using the BLaTM assay and flow cytometry. The selected mutations, identified in α3 (F1019S), β2 (G716V) and β3 (L744P and T746P) subunits, are associated with interstitial lung, nephrotic syndrome and epidermis bullosa (ILNEB), leukocyte adhesion deficiency type I (LAD-I) and Glanzmann thrombasthenia-like syndrome (GT-like), respectively. Our analyses revealed that each mutation impaired the α/β TMD interaction and reduced surface expression of the corresponding integrin when co-expressed with its heterodimeric partner (Figure 6), demonstrating that disrupted α/β TMD interactions underlies integrin loss-of-function disease in humans.

**FIGURE 6.**
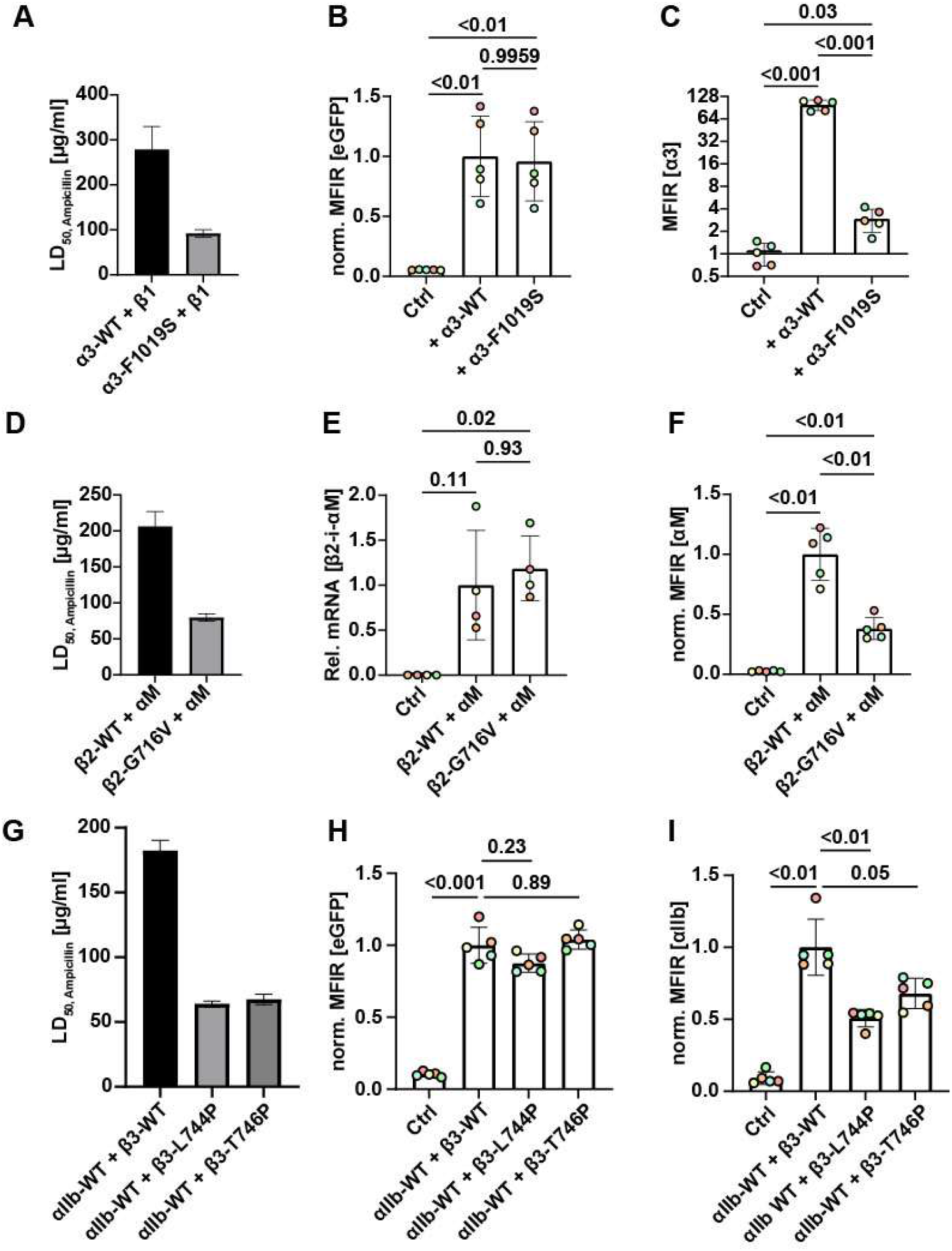
Disease-associated mutations in the TMD of different β-integrin classes impair TMD-TMD association and surface expression. (A,D,G) BlaTM-based measurements of the heterotypic TMD interactions in *E.coli* between indicated wild-type or mutant integrin TMDs fused to N-terminal and C-terminal Bla fragments, respectively. Bacterial survival is indicated as LD50 obtained from global fits of pooled OD600 values from an ampicillin titration series of n=6 biological replicates. (B,C) Flow cytometry-based MFIR measurements of GFP and indicated WT or mutant α3-subunits (detected with the ASC-1 antibody) expression in K562 transduced with a retroviral vector using a 2A self-cleaving peptide (α3-2A-GFP). The MFIR of β1 and α3 is normalized to α3-WT cells. Statistical analysis was done using repeated measures ANOVA with Dunnett’s post-hoc test. Data are shown as mean±SD. (E) Quantitative RT-PCR of indicated WT and mutant β2- and αM-subunit mRNAs in CHO cells using an IRES2 co-expression system (β2-WT-IRES2-αM and β2-G716V-IRES2-αM). Integrin subunit mRNA levels are normalized to actin. (F) Flow cytometry-based MFIR measurements of αM surface expression in β2-WT and β2-G716V cell lines. MFIR is normalized to β2-WT cells. Statistical analysis was performed using Brown-Forsythe and Welch ANOVA with Dunnett T3’s post-hoc test. Data are shown as mean±SD. (H,I) Flow cytometry-based MFIR measurements of GFP, indicated WT or mutant β3 subunits, and αIIb-subunit levels in fibroblasts retrovirally transduced with a T2A and P2A self-cleaving peptides (αIIb-WT-T2A-GFP-P2A-β3-WT; αIIb-WT-T2A-GFP-P2A-β3-L744P; αIIb-WT-T2A-GFP-P2A-β3-T746P). MFIR is normalized to αIIbβ3-WT cells. Statistical analyses were performed using Brown-Forsythe and Welch ANOVA with Dunnett T3’s post-hoc test. Data are shown as mean±SD.

## Discussion

In the ER, newly synthesized α- and β-subunits adopt extended ectodomain conformations before pairing into bent, inactive heterodimers. The bent heterodimers subsequently traffic through the Golgi apparatus to the plasma membrane. At the plasma membrane, reversible and highly dynamic α/β-TMD unclasping and re-clasping, coupled with gross conformational changes in the ectodomain, enable and regulate sustained integrin-ligand binding [25, 27]. In this study, we uncovered a novel and essential role for the TMDs in the ER (Figure 7): they promote the association of newly synthesized, monomeric α- and β-subunits, which brings both subunits in close proximity and thereby facilitates assembly and folding of the α/β-head domain. Thus, integrin subunit TMDs serve a dual role; first they facilitate integrin assembly and folding in the ER, and then they regulate conformational switches at the plasma membrane.

**FIGURE 7.**
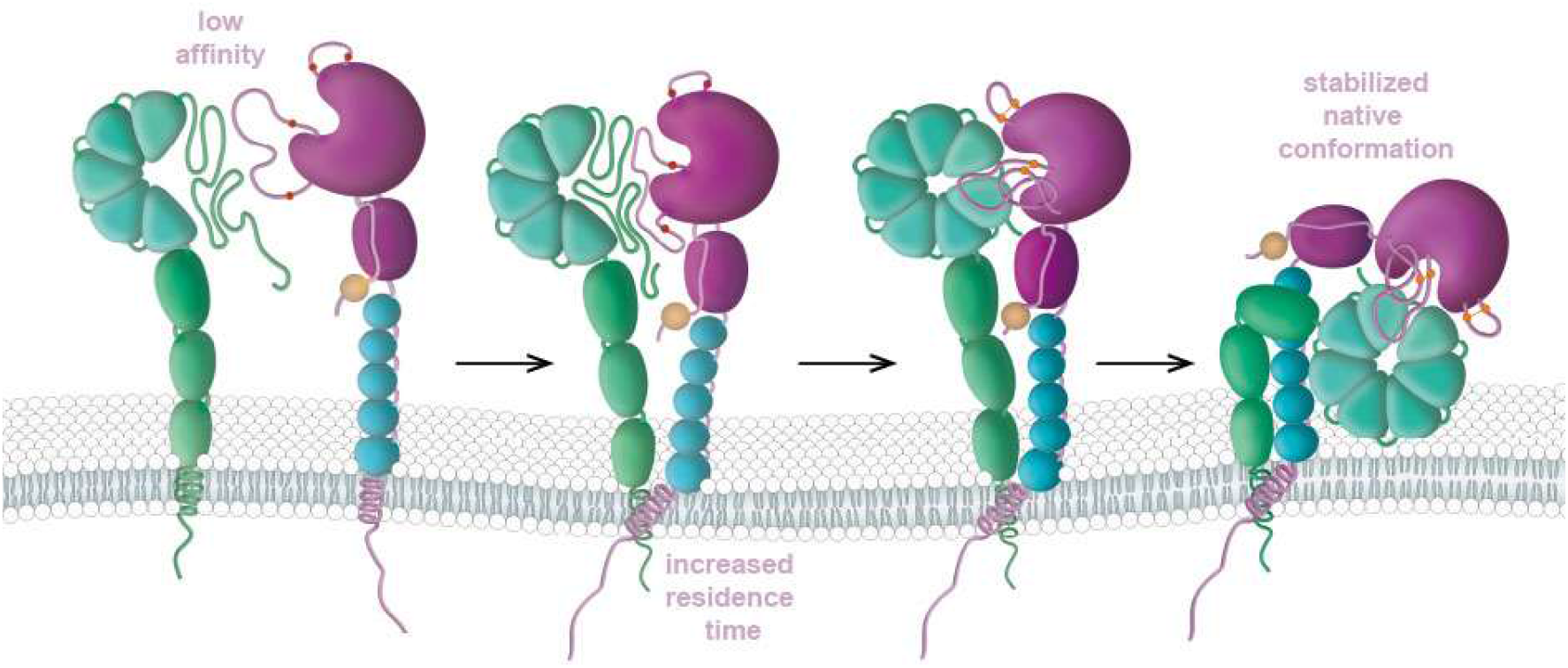
Model of integrin folding in the ER. Upon synthesis, the α and β integrin subunits, interact via their head domains, promoting folding and formation of stable heterodimers that are routed through the secretory pathway. Association of the αTMD-βTMD reinforces the otherwise weak head domain interaction and together with chaperones and divalent cations, facilitates folding and oxidation of critical disulfide bonds in the βI domain to stabilize the heterodimer. Once head domain folding is complete, the ectodomain adopts a bent and is exported to the cell surface.

The initial goal of our study was to determine when and how an elevated fraction of ‘active’ β1 integrins affects murine development. Using genetic engineering, we introduced previously characterized activating mutations from the β3-subunit TMD (G708N/L and L712R) [32,34,35] into the corresponding positions of the β1-subunit TMD (G744N/L and L748R). Contrary to expectations, mice heterozygous for the activating β1 substitutions did not exhibit a dominant gain-of-function phenotype. At homozygosity, β1-LR embryos were severely malformed by E7.5 and underwent resorption, whereas homozygous β1-GN and β1-GL mice appeared phenotypically normal. This defect was not due to reduced protein expression, but rather to ER retention, which was mild in β1-GN and β1-GL and severe in β1-LR. This difference likely reflects the distinct effects of the mutations on the β1-TMD. While the β1-GN and β1-GL mutations subtly alter the local TMD packing, the β1-LR substitution introduces a positively charged residue into the hydrophobic core of the β1-TMD. As previously suggested for β3, the introduction of this charge may shorten the helix, altering its alignment with the α5 TMD within the membrane and potentially destabilizing the entire α-β TMD clasp [35]. These findings demonstrate that activating mutations in the β1-TMD unexpectedly result in a recessive loss-of-function phenotype in mice and reveal a previously unrecognized role for α/β-TMD interaction in preventing β1-integrin retention in the ER.

Our findings indicate that the α/β-TMD association promotes assembly of the non-native, ER-luminal α/β-head domain and correct folding of the βI-domain and β-propeller. In the case of the TMD-disrupting β1-LR substitution, a series of experiments demonstrated impaired formation of α/β-heterodimers and misfolding of the βI-domain. Native gel analysis as well as co-IP followed by MS revealed a pronounced defect in α/β-heterodimerization. Additional evidence for βI-domain unfolding comes from studies with conformation-specific βI-domain-binding antibodies, substitution of the βI-domain with an autonomously folding αI-domain, and impaired disulfide bond formation between C261-C301. Notably, incomplete C261-C301 disulfide bond formation was also observed in β1-WT subunits - which is produced in excess over α-subunits - suggesting that unpaired β1-WT subunits follow the same folding trajectories in the ER as β1-LR subunits.

The analysis of the oxidation state revealed incomplete formation of both, the C207-C213 and C261-C301 disulfide bonds in β1-LR and, to a lesser extent, in the β1-WT subunit. The disulfide bond between C207 and C213 restricts conformational flexibility of the 25-residue specificity-determining loop (SDL) within the integrin βI domain, which promotes an orientation that favors selective ligand recognition [41]. Notably, these cysteine residues are absent in the β4-subunit, indicating that this disulfide bond is not strictly required for βI-domain folding (Figure S4A-C). In contrast, the C261-C301 disulfide bond is conserved across all β-subunits, including β4. This bond stabilizes a rod-shaped protrusion called arginine/lysine finger, which inserts into a groove on the β-propeller of the α-subunit and an adjacent, normally buried, highly hydrophobic β-sheet of the βI-domain (Figure 7). Correct positioning and stabilization of the arginine/lysine finger by C261-C301 is presumably essential for α/β heterodimerization, as mutations of the key arginine or lysine residues in the β3- and β2-subunits, respectively, disrupt α/β-subunit interaction [29,58], consistent with our observation of an almost complete loss of β1 integrins from the cell surface. Furthermore, mutations affecting the corresponding cysteines in the β2 and β4 subunits have been identified in patients with LAD-I and junctional EB, respectively [59,60].

Interestingly, the difference in the abundance of free cysteines between β1-WT or β1-LR is smaller than the difference in mature-to-immature integrin ratio at the cell surface. Several factors may account for this discrepancy. First, misfolding of the βI-domain can involve either of its two disulfide bonds (C207-C213 or C261-C301). Thus, whether these events occur independently or are coupled, their combined contribution to the unfolded fraction would be greater than that of either event alone. Second, integrins can form mixed disulfides with protein disulfide isomerases (PDIs) as folding intermediates [5,8,61], causing these species to be scored as oxidized and thereby diminishing the apparent difference between the two proteins. The effect appears particularly pronounced for β1-LR, as our proteomic data indicate that it engages more extensively with the PDIs, P4hb and Pdia3 and Pdia6.

Although TMD lesions are generally recognized by the ERAD-M pathway [62], our results indicate that the degradation of β1-LR does not occur through this pathway. Chimeric proteins in which the β1-LR-TMD was fused to the autonomously folding extracellular domain of IL2Rα, or in which β1-LR was rescued by fusion with the α5-ectodomain at its N-terminus were efficiently transported to the cell surface. Together, these findings indicate that the β1-LR TMD itself is not sufficient to trigger ERAD-M. Moreover, four ERAD-L proteins (Sel1L-Hrd1 complex, Os9 and Erlec1 adaptors) co-precipitated with both β1-WT and β1-LR. These findings suggest that ERAD-L recognizes both over-produced β1-WT and misfolded ER-retained β1-LR and mediates their retro-translocation across the ER membrane followed by polyubiquitination and proteasomal degradation.

The comparable association of the Sel1L-Hrd1 complex and the ER lectins, Erlec1 and Os9 with β1-WT and β1-LR further suggests that ERAD-L flux of β1 in β1-LR expressing cells does not increase proportionally with the severity of the folding defect leading to the accumulation of β1-LR within the ER. This accumulation, maintained by increased recruitment of chaperones, may increase the frequency of stochastic collisions between α-subunits and β1-LR, thereby promoting productive folding, stable formation of α/β1-LR heterodimers and their subsequent transport to the cell surface, while simultaneously reducing the substrate burden on ERAD. This hypothesis is further supported by the expression of the non-ubiquitinable β1-LR^8xKR^ mutant, which increases the number of possible collision events and consequently β1-LR^8xKR^ surface levels.

Our experiments suggest that integrin heterodimerization is coupled to folding and maturation of the α- and β-subunit head domains. This is consistent with previous studies of αLβ2 biosynthesis demonstrating that folding of the βI-domain of the β2-subunit requires association with αL, whereas folding of the β-propeller domain of the αL-subunit depends on the association with β2 [63,64]. Our findings extend these observations by showing that proper association of the TMDs also contributes to the folding trajectory of the integrin headpiece.

Because regulation of the integrin inactivation/activation cycle is essential for numerous cellular processes, disease-associated mutations that disrupt the α/β-TMD interaction have been implicated in disease pathogenesis by promoting excessive integrin activation without accounting for the potential contribution of reduced surface expression [65–67]. Our analyses of selected TMD point mutations in α3-(associated with ILNEB), β2-(associated with LAD-I) and β3-(associated with GTLS) subunits confirmed impaired TMD interactions and decreased surface levels of integrins. Together with the conserved presence of the two cysteine residues across all known β subunits, these findings point to a principal role for the α/β-TMD interaction in folding of the integrin head domain. The importance of TMD interactions as a quality control checkpoint for ensuring correct folding and surface expression is a recurring phenomenon. For instance, ionic TMD interactions between TCRα and TCRβ subunits and the CD3 complex are essential for both, assembly and surface expression of the mature TCR. Subunits that fail to assemble properly are retro-translocated across the ER membrane into the cytosol and degraded by the proteasome [68–70]. Many polytopic membrane proteins similarly depend on precise packing of their TMDs, with ionic residues often guiding subunit association and subsequent quality control [62,71–74]. However, these static TMD-interactions are incompatible with the reversible and highly dynamic TMD association that regulates integrin activation at the plasma membrane. As overproduced β1-WT subunits are eliminated by the same process, it thus appears, that integrins have co-opted TMD association coupled with assembly-induced folding [75] - which eventually is secured by late disulfide formation - to generate a novel function-surveilling mode of ER quality control. This mechanism ensures that integrins engage the ancient, evolutionary pre-existing ERAD-L machinery to monitor and eliminate unpaired subunits, thereby allowing only properly folded and ‘activatable’ α/β heterodimers to reach the cell surface. This might exclude the non-canonical integrins β4 and β8, for which other activation/ligation mechanisms exist.

Most integrins thus appear to have evolved to fold efficiently only when they encode sufficient TMD interaction strength, thereby licensing them as intrinsically ‘activatable’. This requirement safeguards integrins from dysregulation caused by heterozygous, inherited TMD mutations and thereby neutralizes such potential deleterious alleles. Thus, quantity and quality control of integrins - which includes activation - is secured at the genetic level.

## Material and Methods

### Plasmids and constructs

BlaTM plasmids were generated by Gibson cloning and primers coding for the whole respective TMDs and mutants thereof. A β1-ecto-eGFP-WT construct was generated using the Gibson Assembly® method according to a published construct [38]. Point mutations that generate amino acid changes in β1 (β1-L748R, β1-G744N, β1-G744L and α5-L979D) were introduced in the cDNA using site-directed mutagenesis. The IL2R ectodomain fusion with the β1-WT or β1-LR (IL2R-β1-WT/LR), the exchange of the βI-domain with α2I-domain (α2I:β1-WT/LR), the fusion with the headpiece (α5-SG/2A-β1-WT/LR and the β1-WT/LR-CC/SS and α5-LD/β1-LR- and α5^4xKR^β1-LR^8xKR^-constructs were generated by Gibson cloning. Retroviral vectors containing the integrin subunit cDNAs driven by the EF1α promoter were adapted from the HSC1-GiP self-inactivated retroviral vector (a gift from James Ellis, RRID: Addgene_58254) [76] by replacing either the EGF gene or the puromycin-resistance gene of its bicistronic EGFP-IRES-puromycin cassette with sequences for respective integrin subunits. Alternatively, the entire EGFP-IRES-puromycin cassette was replaced with α5-T2A-β1 or αIIb-T2A-GFP-P2A-β3 or α3-T2A-eGFP sequences. The α5-GFP (a gift from Rick Horwitz (https://www.addgene.org/30450/) was inserted into pRetroQ-N1 vector using restriction cloning.

CRISPR-Cas9-mediated gene deletions in cells were generated using the sgRNA expressing plasmid pSpCas9(BB)-2A-Puro (PX459) V2.0 (a gift from Feng Zhang, RRID: Addgene_62988). The sgRNAs were inserted into PX459 following published protocols [77].

### BlaTM assay

The BlaTM assay was performed following published protocols [36]. Briefly, competent *E. coli* JM83 were co-transformed using the N- and C-Bla plasmids fused to a TMD, plated on LB agar plates containing kanamycin (34 µg/ml) and chloramphenicol (34 µg/ml) (LB_kana/cam_-medium), incubated at 37°C overnight and stored at 4°C for use within 2-3 days. The day prior to the experiment, ∼10 colonies were picked and inoculated in 5 ml LB_kana/cam_-medium. On the day of the experiment, 500 µl of the bacterial suspension was mixed with 4.5 ml of LB_kana/cam_-medium +/- arabinose (inducer) and Isopropyl β-D-1-thiogalactopyranoside (IPTG, negatively regulates induction level) and incubated for 4 h at 37°C on a rotary wheel to induce expression of the fusion protein. Induced samples were adjusted to an OD_600_ of 0.2 and diluted 1:1 with induction medium and grown in 12-well plates containing induction medium supplemented with decreasing concentrations of freshly prepared ampicillin for 19 hours at 37°C. The next day, 200 µl of bacterial solution were transferred to a 96-well plate and OD_600_ was measured for calculating the LD_50_.

For GFP expression controls, induced and non-induced samples were collected and 200 µl were transferred to a 96-well plate, centrifuged and resuspended in PBS. Raw GFP expression was measured in a BioTek Synergy H1 Multimode Reader (λ_ex_ = 485 nm, λ_em_ = 520 nm).

### Generation of transgenic animals

Mice carrying the TMD germline mutations followed the same strategy as described previously [37]. Briefly, targeting constructs were generated by restriction cloning and site directed mutagenesis of mouse genomic clones [78]. The final 16 kb targeting constructs, which contained within intron 14 a loxP site, a truncating exon 15 followed by a stop codon, a neomycin resistance cassette. a second loxP site and exon 15 carrying the point mutation, were electroporated into R1 ES cells. After selection, recombination-positive ES cell clones were identified by Southern blotting of BamHI-digested genomic DNA using 5’- and 3’-external probes and injected into C57B6 blastocysts [37]. After germline transmission, the truncated exon 15 and neomycin cassette were removed by breeding to deleter-Cre strain [79] leading to β1^wt/G744N^, β1^wt/G744L^, β1^wt/G748R^ mice for subsequent homozygous interbreeding. Genotyping was performed on genomic DNA from tail biopsies or hair follicles by PCR.

### Cell lines

All cell lines were cultured at 37°C and 5% CO_2_ and routinely tested for mycoplasma contamination. Fibroblasts were maintained in DMEM supplemented with 10% FBS. K562 cells were cultured in RPMI-1640 medium supplemented with 10% FBS, and CHO cells in Ham’s F12 medium supplemented with GlutaMAX and 10% FBS. Generation and culture of β1^flox/flox^ and β1^-/-^ cells were described previously [44]. Fibroblasts with null mutations in α5 as well as β1 integrin genes (α5^-/-^β1^-/-^; DKO) were generated from SV40-immortalized β1^flox/flox^ adult kidney fibroblasts by CRISPR-Cas9-mediated deletion of the gene encoding the α5 subunit using two different sgRNAs (see above) and two rounds of FACS sorting for the α5-negative cell population.

Subsequently, the α5-deficient β1^flox/flox^-fibroblasts were transduced with an adenovirus expressing Cre-GFP and sorted for GFP^high^-β1^low^ 48h later. The resulting cell population was then re-sorted for β1^negative^ cells, and seeded at a density of 100 cells per well in a 24-well plate. Loss of α5 and β1 expression was verified by flow cytometry and Western blotting.

Stable re-expression of the wild-type and mutant α5 and β1 subunits in α5^-/-^β1^-/-^ DKO fibroblasts was achieved by retroviral transduction. The transduced cells were enriched either by FACS or puromycin selection using 2.5-10 µg/ml of puromycin for 2-4 days.

### Blue native (BN) PAGE

Around 150-200.000 cells were seeded in a 6-well plate one day prior to the experiment. The next day, cells were washed once with PBS and lysed in ice-cold 100 µl BN-lysis buffer (20 mM Bis-Tris-HCl, pH 7.5, 750 mM ε-caproic acid, 1% digitonin). Lysates were collected by scraping and cleared by centrifugation at 16.000 x *g* for 10 min at 4°C. Protein concentrations were determined using the BCA assay and adjusted for equal amounts. The lysates were then mixed with an equal volume of Native PAGE loading buffer (50% Native PAGE sample buffer and 4% G-250) and 16 μl of the protein samples were loaded into each well of a 15-well 4-16% gradient gel (Bis-Tris, Invitrogen). Samples were run on the gel according to the manufacturer’s instructions using the appropriate dark blue or light blue cathode buffer systems. The gels were rinsed in 10% ethanol for 10 minutes, washed once with Milli-Q^®^ grade water after which the proteins were denatured using 10x SDS running buffer supplemented with 0.4% β-mercaptoethanol for 1h at RT. Subsequently, gels were washed three times with Milli-Q^®^ grade water and subjected to standard wet transfer conditions for Western blotting.

### Endo H and PNGase F digestion

Protein lysates were prepared in lysis buffer containing 1% Triton X-100, 0.1% sodium deoxycholate, 150 mM NaCl, cOmplete protease inhibitor cocktail, and 50 mM Tris-HCl (pH 7.5). For deglycosylation, 10 μg of protein lysate was denatured at 100°C for 10 min and then incubated with either Endo H (P0702S, NEB) or PNGase F (P0704S, NEB) at 37°C for 1 h, according to the manufacturer’s instructions. Samples were subsequently analyzed by 8% SDS-PAGE.

### Immunoprecipitation

Antibody binding to mature β1 and its precursor were analyzed by immunoprecipitation. Briefly, protein A/G beads were equilibrated by washing with β1-lysis buffer (1% Triton X-100, 0.1% sodium deoxycholate, 150 mM NaCl, cOmplete protease inhibitor, 50 mM Tris/HCl, pH 7.5), allowed to bind antibodies (2 µg/30 µl beads) and washed three times with β1-lysis buffer. The following antibodies were used: βI-domain specific mAbs TS2/16, P4C10, mAb13, 12G10, hybrid domain specific mAbs Huts4, Huts21 and leg/PSI-domain specific antibodies 9EG7, N29 and 8E3. Cells were washed once with cold PBS, scraped in β1-lysis buffer, collected and sonicated on ice for 10 seconds and centrifuged at 16.000 x *g* for 10 min. Cleared protein lysates (1 mg) were added to 30 µl of the antibody-bound beads. incubated overnight on a rotary wheel at 4°C and washed 3-times with β1-lysis buffer. The proteins were eluted from the protein A/G beads by boiling in 4x SDS sample buffer containing 2-mercaptoethanol, separated by SDS-PAGE and Western blotted.

Immunoprecipitation of β1-ecto-GFP from cells treated with 5 µM CB-5083 or DMSO for 6 h was carried out as described above, except that the cells were lysed in hypotonic lysis buffer (10 mM HEPES, 10 mM KCl, 1% CHAPS, complete protease inhibitor, pH 7.9) and anti-GFP (nanobody) magnetic beads (30 µl) were used to immunoprecipitate β1-ecto-GFP from 2 mg of protein lysate.

### Membrane fractionation

Membrane (pellet) and cytosolic (supernatant) extracts were generated as follows [80]. Cells at 80-90% confluency in 10 cm dishes were washed once with PBS and once with a hypotonic salt solution (100 mM NaCl). The cells were subjected to either sodium carbonate extraction buffer (100 mM sodium carbonate, pH 11.5, ice-cold) or high salt extraction buffer (10 mM Tris-HCl, 500 mM NaCl, pH 8.0). Extracts were then extruded 10 times through a 27G needle and centrifuged at 150.000 x *g* for 1h at 4°C. After dissolving the pellets in respective buffers, the protein suspensions and supernatants were mixed with an equal volume of 4x SDS sample buffer, boiled and Western blotted.

### Flow cytometry

Cells were detached using PBS-EDTA (5 mM EDTA in PBS), counted and seeded the day prior to the experiment at a cell density that result in 70-80% confluency after 24 h (500-750.000 cells/10 cm dish). On the day of the experiment, cells were detached with PBS-EDTA and approximately 200.000 cells were incubated on ice for 30 min with primary antibodies in 100 µl FACS buffer (3% BSA, 5 mM EDTA in PBS). The cells were washed once with cold PBS, centrifuged and incubated with A647-conjugated secondary antibodies for 30 min at 4 °C, or analyzed directly. After washing, secondary antibody binding was analyzed by flow cytometry using the LSR Fortessa X-20 Cell Analyzer. For quantification, geometric mean fluorescence intensities of cellular populations were calculated using FlowJo 10.6.1.

### Immunofluorescence

For immunofluorescence staining, cells were seeded on FN (10 µg/ml) or serum vitronectin-coated (for endogenous fibronectin staining) ibidi µ-slides and allowed to adhere and spread 6h. After washing the wells with 1 mM Ca^2+^/Mg^2+^-supplemented with PBS, cells were fixed with 4% PFA in PBS for 5 or 10 min, washed, permeabilized with 0.1% Triton X-100 in PBS for 10 min, blocked with 3% BSA in PBS supplemented with 0.1% Tween-20 and incubated with primary and then secondary antibodies (in the blocking buffer) for 2h at room temperature or overnight at 4°C. After the incubation, the cells were washed 4 times for 5 minutes with PBS. DAPI (1:10.000) was added in the first washing step. Cells were embedded in ibidi mounting medium and imaged or stored at 4°C. Samples were imaged using a Zeiss LSM780 or LSM980 confocal laser scanning microscope using ZEN software. Images were analyzed with Fiji ImageJ software.

For immunohistochemistry embryos at embryonic day (E) 7.5 were dissected and processed as described previously [81]. Briefly, harvested embryos were fixed in 4% PFA for 2 h at 4°C, followed by 3 washes with cold PBS, dehydrated with 70% and 100% ethanol, cleared in xylene and embedded in paraffin wax. Sections of 5 µm thickness from the paraffin wax were permeabilized with 0.1% Triton X-100 (v/v) in PBS for 15 min and then blocked in 3% BSA for 2 h at RT followed by incubation with the primary antibody overnight at 4°C and secondary for 2 h at RT. Images were collected on a Zeiss (Jena) LSM780 confocal laser scanning microscope.

### Metabolic labeling

For metabolic labeling, 1.5×10^6^ cells were seeded on FN and grown to 80% confluency overnight. Cells were incubated 45 min in methionine/cysteine-free medium supplemented with 10% dialyzed FBS (labeling medium) and indicated chemicals. For labeling, cells were incubated in labeling medium supplemented with additional [^35^S]methionine/cysteine (250 µCi / 10 cm plate) and indicated inhibitors or DMSO for 30 min. After labeling, cells were washed twice with complete PBS and either lysed directly (0h) or chased for 6h in DMEM supplemented with 10% FBS. Cells were lysed in β1-lysis buffer, collected and cleared by centrifugation. The lysates were incubated with a homemade polyclonal anti-β1 antibody-protein A/G bead conjugate for 2h at 4°C [82]. After removal of lysates and washing of beads, proteins were eluted by boiling in 4x Laemmli buffer. Samples were subjected to SDS-PAGE, gels were fixed and dried and exposed to phosphor screens. Images were collected on a Typhoon^TM^ system.

### VISTA proteomics

For proteomic analysis of β1 interactors in the ER [83], 3×10^6^ cells were seeded in 15 cm dishes the day prior to the experiment. Cells were treated with 5 µM CB-5083 for 6h and lysed in hypotonic lysis buffer, followed by scraping, sonication and centrifugation. Protein lysates (2 mg) were added to 30 µl of anti-GFP nanobeads, incubated overnight on a rotary wheel at 4°C and washed three times with the hypotonic lysis buffer, followed by two washes with PBS to remove excess detergent. Beads were incubated with 50 µl of SDC buffer (1% SDC, 40 mM 2-chloroacetamide, 10 mM tris(2-carboxyethyl)-phosphine (TCEP), 100 mM Tris, pH 8.0) at 37 °C. Following a 20 min incubation at 37 °C, the samples were diluted with 50 µl of MS grade water (VWR). Proteins were then digested overnight at 37 °C by adding 0.5 µg trypsin (Promega) or GluC (Promega). The resulting supernatant was collected, beads were washed with 100 µL of buffer A (0.1% formic acid), and unified supernatants were acidified with trifluoroacetic acid (TFA; Merck) to a final concentration of 1%. Precipitated SDC was removed via centrifugation, and the peptide mixture was desalted using SCX StageTips. After elution, peptides were either vacuum dried and dissolved in buffer A (0.1% formic acid).

Desalted peptides were separated at a flow rate of 350 µl/min on a C18 column (30 cm, inner diameter: 75 microns; packed in-house with ReproSil-Pur C18-AQ 1.9-micron beads, Dr. Maisch GmbH) via the Easy-nLC 1200 (Thermo Fisher Scientific) using a gradient of buffer A (0.1% formic acid) to buffer B (80% acetonitrile, 0.1% formic acid). The composition of buffer B was gradually increased: initially set at 8%, it was ramped up to 30% over 35 min. Subsequently, there was an increase to 35% over 10 min, followed by a further increase to 58% over the next 5 min. Finally, the buffer was adjusted to 95% over the subsequent 5 min and maintained at this concentration for another 5 min. The LC-system was coupled to the timsTOF Pro mass spectrometer (Bruker Daltonics), and data acquisition was performed using otofControl 6.0. The mass spectrometer was operated in data-dependent PASEF mode with one survey TIMS-MS and ten PASEF MS/MS scans per acquisition cycle. Analysis was performed in a mass scan range from 100-1700 m/z and an ion mobility range from 1/K0 = 1.6 Vs cm-2 to 0.6 Vs cm-2 using equal ion accumulation and ramp time in the dual TIMS analyzer of 100 ms each at a spectra rate of 9.52 Hz. Suitable precursor ions for MS/MS analysis were isolated in a window of 2 Th for m/z < 700 and 3 Th for m/z > 700 by rapidly switching the quadrupole position in sync with the elution of precursors from the TIMS device. The collision energy was lowered stepwise as a function of increasing ion mobility, starting from 52 eV for 0-19% of the TIMS ramp time, 47 eV from 19-38%, 42 eV from 38-57%, 37 eV from 57-76%, and 32 eV until the end. We made use of the m/z and ion mobility information to exclude singly charged precursor ions with a polygon filter mask and further used ‘dynamic exclusion’ to avoid re-sequencing of precursors that reached a ‘target value’ of 20,000 a.u. The ion mobility dimension was calibrated linearly using three ions from the Agilent ESI LC/MS tuning mix (m/z, 1/K0: 622.0289, 0.9848 Vs cm-2; 922.0097, 1.1895 Vs cm-2; 1221.9906, 1.3820 Vs cm-2).

The recorded data underwent processing using the MaxQuant computational platform (version 1.6.17.0). In brief, the peak list was searched against the Mouse and Human Uniprot database (downloaded in 2020). Cysteine carbamidomethylation was set as static modification, and methionine oxidation, deamidation, lysine-biotinylation and N-terminal acetylation as variable modifications. Trypsin or GluC were set as protease. The match-between-run option was enabled, and proteins were quantified across samples using the label-free quantification algorithm in MaxQuant generating label-free quantification (LFQ) intensities. Intensities of identified proteins were loaded into Perseus software (version 1.6.2.3). Doublet entries stemming from different enzymes (columns) or Protein IDs (rows) were summed. Data were log-transformed, missing values imputed from a normal distribution, and filtered to remove reverse identifications and proteins with fewer than three unique peptides.

### Cysteine oxidation mapping

2×10^6^ cells (β1-KO fibroblasts and β1-KO fibroblasts expressing either mouse or human β1-WT or β1-LR, respectively) were seeded in 16 15-cm culture dishes one day prior to the experiment. Dishes were organized in random quadruplicates and washed once with complete PBS supplemented with 40 mM N-Ethylmaleimide (NEM) and put on ice. Subsequently, cells were lysed in 1.0 ml β1-lysis buffer supplemented with 40 mM NEM, scraped, sonicated and centrifuged. The supernatants (4 mg protein) were added to 35 µl of protein A/G beads, pre-incubated with 2 µg of the anti-β1 mAb JB1A or a homemade anti-mouse β1 serum, and incubated overnight at 4°C. The next day, the beads were washed three times with β1-lysis buffer and twice with PBS. Beads were incubated with 50 µl of PBS supplemented with 10 mM of TCEP for 15 min at 37 °C, then 25 µl of *d5*-NEM (10 mM stock solution) was added and incubated for an additional 15 min. After the protein-containing samples were diluted with 75 µl of MilliQ water, they were treated with 0.5 µg sequencing grade Trypsin (Promega) or Glu-C (Promega) at 37 °C overnight. The samples were then centrifuged and the supernatant collected in a fresh tube. The beads were washed with 0.1% formic acid, centrifuged and the supernatant mixed with the supernatant previously centrifuged. The mixed supernatants with the digested peptides were then acidified with trifluoroacetic acid (TFA) to a final concentration of 1% and loaded onto Evotips. Evotips were eluted onto a 15-cm column (PepSep C18 15cmx150µm, 1.5 µm, Bruker Daltonics) via the Evosep One HPLC system (Evosep). The column was heated to 50°C and peptides were separated using the “30 samples per day” method. Using the nano-electrospray interface, eluting peptides were directly sprayed onto the timsTOF Pro mass spectrometer (Bruker Daltonics) and data acquisition was performed using timsControl. The MS was operated in data-independent (DIA) PASEF mode. Analysis was performed in a mass scan range from 100-1700 m/z and an ion mobility range from 1/K0 = 0.70 Vs cm-2 to 1.30 Vs cm-2 using equal ion accumulation and ramp time in the dual TIMS analyzer of 100 ms each at a spectra rate of 9.52 Hz. diaPASEF scans were acquired a mass scan range from 350.2-1199.9 Da and an ion mobility range from 1/K0 = 0.70 Vs cm-2 to 1.30 Vs cm-2. Collision energy was ramped linearly as a function of the mobility from 45 eV at 1/K0 = 1.30 Vs cm−2 to 27 eV at 1/K0 = 0.85 Vs cm−2. In complete, 42 diaPASEF windows were distributed to one TIMS scan each at switching Th precursor isolation windows which led to an estimated cycle time of 2.21 s.

The ion mobility dimension was calibrated linearly using three ions from the Agilent ESI LC/MS tuning mix (m/z, 1/K0: 622.0289, 0.9848 Vs cm-2; 922.0097, 1.1895 Vs cm-2; 1221.9906, 1.3820 Vs cm2).

### Data analysis of cysteine oxidation experiments

Raw data was processed using the Spectronaut® software (18.0). For GluC and trypsin runs peak lists were searched against a minimized FASTA-file containing the 150 highest intensity proteins (additionally including human ITGB1 and mouse Pdia2/4/5) of a pre-search using the whole mouse genome (downloaded from Uniprot in July 2023). Light and heavy NEM were set as “Label” (+125/130 Da, label-workflow), allowing for up to 7 modifications/peptide, two missed cleavages and methionine oxidation as variable modification.

Obtained peptide lists for GluC and tryptic peptides were combined and filtered for β1 peptides with cysteine labels. Cysteine labeled β1 peptides were filtered for entries containing less than 6 identifications with intensities > 10000 in the MS light channel for the β1-KO control samples. For a given cysteine residue, intensities (Int) of technical replicates (j) and individual peptides (i) were summed up to result in a summed intensity (SI) for the respective label channel and quantification method (MS1 or MS2):

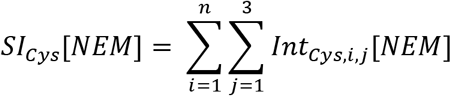

Fractions of free cysteines were calculated for every individual cysteine as average of the fractions obtained from MS and MS/MS measurements:

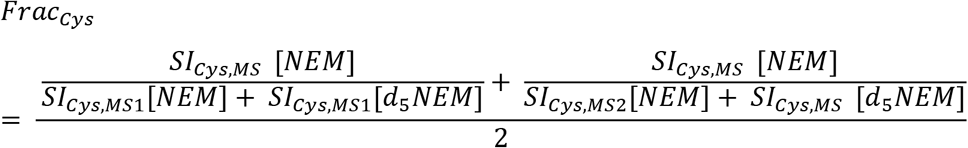

Obtained values of biological replicates for β1-WT/LR samples of individual cysteines were aggregated and analyzed in GraphPad software using multiple t-tests.

### Statistical analysis

Statistical analysis was done primarily using GraphPad Prism version 9 or 10. If not mentioned otherwise, ANOVA was used and values depicted are averages ± standard deviation with multiple t-test corrected p-values above the graph. For the BLaTM assay, OD_600_-values of *E. coli* suspension cultures of ampicillin gradient arrays were aggregated for all replicates and plotted in OriginPro calculating non-linear dose response curves using the Levenberg-Marquardt algorithm (no weighting or filtering).

## Acknowledgements

We thank Michal Grzejszczyk, Hannelore Reiter, Simone Bach, Nicole Krombholz and Anja Wehner for technical assistance, Drs. Martin Spitaler and Giovanni Cardone from the Imaging Facility, and Soo Jin Min-Weißenhorn from the animal house for technical support. We thank Dr. Jieqing Zhu for providing the *in silico* predicted structure of α5β1. This work was supported by the European Research Council (ERC) under the European Union’s Horizon 2020 research and innovation program (grant agreement No. 810104 – Point), the Max Planck Society (to RF), the National Natural Science Foundation of China (NSFC 82670856 to SSG) and the Shenzhen Maternity and Child Healthcare Hospital (KYQD20260003 to SSG) affiliated to the Southern Medical University.

## Author contributions

The project was conceived and initiated by RF, who supervised the study and secured funding, with additional supervision and funding by SSG and RTB. Experiments were designed by JMR, RTB, HM, DP, SSG and RF. Experimental work was carried out by JMR, with additional experiments provided by RTB, DP, HM, SSG, PWK, BS and WQ. DL and AS provided advice, reagents and methodologies. JMR performed the formal analysis and interpretation of the data and developed the mechanistic framework underlying the results, in discussions with RF, AS and SSG. JMR prepared the figures, with contributions from RTB, SSG and WQ. The first draft was written by JMR. The manuscript was finalized by RF, AS, JMR and SSG, and approved by all authors.

## References

[1] J. W. Tamkun, D. W. DeSimone, D. Fonda, R. S. Patel, C. Buck, A. F. Horwitz, R. O. Hynes, Cell 1986, 46 (2), 271, 10.1016/0092-8674(86)90744-0.

[2] I. D. Campbell, M. J. Humphries, Cold Spring Harb Perspect Biol 2011, 3 (3), 10.1101/cshperspect.a004994.

[3] S. K. Akiyama, K. M. Yamada, J Biol Chem 1987, 262 (36), 17536.

[4] J. Heino, R. A. Ignotz, M. E. Hemler, C. Crouse, J. Massague, J Biol Chem 1989, 264 (1), 380.

[5] C. E. Jessop, S. Chakravarthi, N. Garbi, G. J. Hammerling, S. Lovell, N. J. Bulleid, EMBO J 2007, 26 (1), 28, 10.1038/sj.emboj.7601505.

[6] M. Lenter, D. Vestweber, J Biol Chem 1994, 269 (16), 12263.

[7] N. A. Hotchin, A. Gandarillas, F. M. Watt, J Cell Biol 1995, 128 (6), 1209, 10.1083/jcb.128.6.1209.

[8] C. E. Jessop, T. J. Tavender, R. H. Watkins, J. E. Chambers, N. J. Bulleid, J Biol Chem 2009, 284 (4), 2194, 10.1074/jbc.M808054200.

[9] J. Li, M. H. Jo, J. Yan, T. Hall, J. Lee, U. Lopez-Sanchez, S. Yan, T. Ha, T. A. Springer, Cell 2024, 187 (12), 2990, 10.1016/j.cell.2024.04.049.

[10] J. Takagi, B. M. Petre, T. Walz, T. A. Springer, Cell 2002, 110 (5), 599, 10.1016/s0092-8674(02)00935-2.

[11] T. Xiao, J. Takagi, B. S. Coller, J. H. Wang, T. A. Springer, Nature 2004, 432 (7013), 59, 10.1038/nature02976.

[12] A. Chigaev, T. Buranda, D. C. Dwyer, E. R. Prossnitz, L. A. Sklar, Biophys J 2003, 85 (6), 3951, 10.1016/S0006-3495(03)74809-7.

[13] T. I. Moore, J. Aaron, T. L. Chew, T. A. Springer, Cell Rep 2018, 22 (7), 1903, 10.1016/j.celrep.2018.01.062.

[14] T. Kamata, M. Handa, S. Ito, Y. Sato, T. Ohtani, Y. Kawai, Y. Ikeda, S. Aiso, J Biol Chem 2010, 285 (49), 38428, 10.1074/jbc.M110.139667.

[15] M. Kim, C. V. Carman, T. A. Springer, Science 2003, 301 (5640), 1720, 10.1126/science.1084174.

[16] B. H. Luo, T. A. Springer, J. Takagi, PLoS Biol 2004, 2 (6), e153, 10.1371/journal.pbio.0020153.

[17] J. Zhu, C. V. Carman, M. Kim, M. Shimaoka, T. A. Springer, B. H. Luo, Blood 2007, 110 (7), 2475, 10.1182/blood-2007-03-080077.

[18] Z. Sun, M. Costell, R. Fassler, Nat Cell Biol 2019, 21 (1), 25, 10.1038/s41556-018-0234-9.

[19] S. Tadokoro, S. J. Shattil, K. Eto, V. Tai, R. C. Liddington, J. M. de Pereda, M. H. Ginsberg, D. A. Calderwood, Science 2003, 302 (5642), 103, 10.1126/science.1086652.

[20] K. L. Wegener, A. W. Partridge, J. Han, A. R. Pickford, R. C. Liddington, M. H. Ginsberg, I. D. Campbell, Cell 2007, 128 (1), 171, 10.1016/j.cell.2006.10.048.

[21] A. Meves, C. Stremmel, R. T. Bottcher, R. Fassler, J Invest Dermatol 2013, 133 (12), 2722, 10.1038/jid.2013.232.

[22] C. T. Lefort, J. Rossaint, M. Moser, B. G. Petrich, A. Zarbock, S. J. Monkley, D. R. Critchley, M. H. Ginsberg, R. Fassler, K. Ley, Blood 2012, 119 (18), 4275, 10.1182/blood-2011-08-373118.

[23] J. Aretz, M. Aziz, N. Strohmeyer, M. Sattler, R. Fassler, Nat Struct Mol Biol 2023, 30 (12), 1913, 10.1038/s41594-023-01139-9.

[24] M. A. Bodescu, J. Aretz, M. Grison, M. Rief, R. Fassler, Proc Natl Acad Sci U S A 2023, 120 (26), e2218116120, 10.1073/pnas.2218116120.

[25] S. Tiwari, J. A. Askari, M. J. Humphries, N. J. Bulleid, J Cell Sci 2011, 124 (Pt 10), 1672, 10.1242/jcs.084483.

[26] J. Li, Y. Su, W. Xia, Y. Qin, M. J. Humphries, D. Vestweber, C. Cabanas, C. Lu, T. A. Springer, EMBO J 2017, 36 (5), 629, 10.15252/embj.201695803.

[27] J. Zhu, B. H. Luo, P. Barth, J. Schonbrun, D. Baker, T. A. Springer, Mol Cell 2009, 34 (2), 234, 10.1016/j.molcel.2009.02.022.

[28] W. B. Mitchell, J. Li, M. Murcia, N. Valentin, P. J. Newman, B. S. Coller, Blood 2007, 109 (9), 3725, 10.1182/blood-2006-11-058420.

[29] J. P. Xiong, T. Stehle, B. Diefenbach, R. Zhang, R. Dunker, D. L. Scott, A. Joachimiak, S. L. Goodman, M. A. Arnaout, Science 2001, 294 (5541), 339, 10.1126/science.1064535.

[30] T. L. Lau, C. Kim, M. H. Ginsberg, T. S. Ulmer, EMBO J 2009, 28 (9), 1351, 10.1038/emboj.2009.63.

[31] B. W. Berger, D. W. Kulp, L. M. Span, J. L. DeGrado, P. C. Billings, A. Senes, J. S. Bennett, W. F. DeGrado, Proc Natl Acad Sci U S A 2010, 107 (2), 703, 10.1073/pnas.0910873107.

[32] B. H. Luo, C. V. Carman, J. Takagi, T. A. Springer, Proc Natl Acad Sci U S A 2005, 102 (10), 3679, 10.1073/pnas.0409440102.

[33] H. Zhu, D. G. Metcalf, C. N. Streu, P. C. Billings, W. F. Degrado, J. S. Bennett, J Mol Biol 2010, 401 (5), 882, 10.1016/j.jmb.2010.06.062.

[34] R. Li, N. Mitra, H. Gratkowski, G. Vilaire, R. Litvinov, C. Nagasami, J. W. Weisel, J. D. Lear, W. F. DeGrado, J. S. Bennett, Science 2003, 300 (5620), 795, 10.1126/science.1079441.

[35] A. W. Partridge, S. Liu, S. Kim, J. U. Bowie, M. H. Ginsberg, J Biol Chem 2005, 280 (8), 7294, 10.1074/jbc.M412701200.

[36] C. Schanzenbach, F. C. Schmidt, P. Breckner, M. G. Teese, D. Langosch, Sci Rep 2017, 7, 43476, 10.1038/srep43476.

[37] A. Czuchra, H. Meyer, K. R. Legate, C. Brakebusch, R. Fassler, J Cell Biol 2006, 174 (6), 889, 10.1083/jcb.200604060.

[38] C. Huet-Calderwood, F. Rivera-Molina, D. V. Iwamoto, E. B. Kromann, D. Toomre, D. A. Calderwood, Nat Commun 2017, 8 (1), 570, 10.1038/s41467-017-00646-w.

[39] H. Zhang, D. S. Zhu, J. Zhu, Comput Struct Biotechnol J 2023, 21, 4497, 10.1016/j.csbj.2023.09.022.

[40] J. Emsley, C. G. Knight, R. W. Farndale, M. J. Barnes, R. C. Liddington, Cell 2000, 101 (1), 47, 10.1016/S0092-8674(00)80622-4.

[41] J. Takagi, D. P. DeBottis, H. P. Erickson, T. A. Springer, Biochemistry 2002, 41 (13), 4339, 10.1021/bi016047u.

[42] M. Nagae, S. Re, E. Mihara, T. Nogi, Y. Sugita, J. Takagi, J Cell Biol 2012, 197 (1), 131, 10.1083/jcb.201111077.

[43] N. Berner, K. R. Reutter, D. H. Wolf, Annu Rev Biochem 2018, 87, 751, 10.1146/annurev-biochem-062917-012749.

[44] R. T. Bottcher, C. Stremmel, A. Meves, H. Meyer, M. Widmaier, H. Y. Tseng, R. Fassler, Nat Cell Biol 2012, 14 (6), 584, 10.1038/ncb2501.

[45] R. M. Dai, C. C. Li, Nat Cell Biol 2001, 3 (8), 740, 10.1038/35087056.

[46] E. Jarosch, C. Taxis, C. Volkwein, J. Bordallo, D. Finley, D. H. Wolf, T. Sommer, Nat Cell Biol 2002, 4 (2), 134, 10.1038/ncb746.

[47] Y. Ye, H. H. Meyer, T. A. Rapoport, Nature 2001, 414 (6864), 652, 10.1038/414652a.

[48] S. Schoebel, W. Mi, A. Stein, S. Ovchinnikov, R. Pavlovicz, F. DiMaio, D. Baker, M. G. Chambers, H. Su, D. Li, T. A. Rapoport, M. Liao, Nature 2017, 548 (7667), 352, 10.1038/nature23314.

[49] R. S. Hegde, H. L. Ploegh, Curr Opin Cell Biol 2010, 22 (4), 437, 10.1016/j.ceb.2010.05.005.

[50] P. M. Kambli, U. A. Bargir, R. M. Yadav, M. R. Gupta, A. D. Dalvi, G. Hule, M. Kelkar, S. Sawant-Desai, P. Setia, N. Jodhawat, N. Nambiar, A. Dhawale, P. Gaikwad, S. Shinde, P. Taur, V. Gowri, A. Pandrowala, A. Gupta, V. Joshi, M. Sharma, K. Arora, R. K. Pilania, H. Chaudhary, A. Agarwal, S. Katiyar, S. Bhattad, S. Ramprakash, R. Cp, A. Jayaram, V. Gornale, R. Raj, R. Uppuluri, M. Sivasankaran, D. Munirathnam, H. P. Lashkari, M. Kalra, A. Sachdeva, A. Sharma, S. Balaji, G. M. Govindraj, S. Karande, R. Nanavati, M. Manglani, G. Subramanyam, A. Sampagar, I. Ck, P. Gutha, S. Kanakia, S. P. Mundada, V. Krishna, S. Nampoothiri, S. Nemani, A. Rawat, M. Desai, M. Madkaikar, Front Immunol 2020, 11, 612703, 10.3389/fimmu.2020.612703.

[51] N. Parvaneh, S. Mamishi, A. Rezaei, N. Rezaei, B. Tamizifar, L. Parvaneh, R. Sherkat, B. Ghalehbaghi, S. Kashef, Z. Chavoshzadeh, A. Isaeian, F. Ashrafi, A. Aghamohammadi, J Clin Immunol 2010, 30 (5), 756, 10.1007/s10875-010-9433-2.

[52] B. Esmaeili, M. Ghadami, M. R. Fazlollahi, S. Niroomanesh, L. Atarod, Z. Chavoshzadeh, Z. Moradi, Z. Alizadeh, Z. Pourpak, Iran J Allergy Asthma Immunol 2014, 13 (1), 61.

[53] C. Ghevaert, A. Salsmann, N. A. Watkins, E. Schaffner-Reckinger, A. Rankin, S. F. Garner, J. Stephens, G. A. Smith, N. Debili, W. Vainchenker, P. G. de Groot, J. A. Huntington, M. Laffan, N. Kieffer, W. H. Ouwehand, Blood 2008, 111 (7), 3407, 10.1182/blood-2007-09-112615.

[54] S. Kunishima, H. Kashiwagi, M. Otsu, N. Takayama, K. Eto, M. Onodera, Y. Miyajima, Y. Takamatsu, J. Suzumiya, K. Matsubara, Y. Tomiyama, H. Saito, Blood 2011, 117 (20), 5479, 10.1182/blood-2010-12-323691.

[55] A. Jayo, I. Conde, P. Lastres, C. Martinez, J. Rivera, V. Vicente, C. Gonzalez-Manchon, Haematologica 2010, 95 (7), 1158, 10.3324/haematol.2009.018572.

[56] Y. Kobayashi, H. Matsui, A. Kanai, M. Tsumura, S. Okada, M. Miki, K. Nakamura, S. Kunishima, T. Inaba, M. Kobayashi, Br J Haematol 2013, 160 (4), 521, 10.1111/bjh.12160.

[57] S. Morais, J. Oliveira, C. Lau, M. Pereira, M. Goncalves, C. Monteiro, A. R. Goncalves, R. Matos, M. Sampaio, E. Cruz, I. Freitas, R. Santos, M. Lima, PLoS One 2020, 15 (12), e0235136, 10.1371/journal.pone.0235136.

[58] V. Gupta, J. L. Alonso, T. Sugimori, M. Essafi, J. P. Xiong, M. A. Arnaout, J Immunol 2008, 180 (3), 1713, 10.4049/jimmunol.180.3.1713.

[59] H. Nawaz Tipu, R. Raza, S. Jaffar, A. Khan, M. Z. Anwar, W. Ahmad, S. I. Raza, Immunobiology 2020, 225 (3), 151938, 10.1016/j.imbio.2020.151938.

[60] L. M. Mariath, J. T. Santin, J. A. Frantz, M. J. R. Doriqui, L. Schuler-Faccini, A. E. Kiszewski, Clin Genet 2021, 99 (1), 29, 10.1111/cge.13792.

[61] D. Bilches Medinas, S. Malik, E. Yildiz-Bolukbasi, J. Borgonovo, M. J. Saaranen, H. Urra, E. Pulgar, M. Afzal, D. Contreras, M. T. Wright, F. Bodaleo, G. Quiroz, P. Rozas, S. Mumtaz, R. Diaz, C. Rozas, F. Cabral-Miranda, R. Pina, V. Valenzuela, O. Uyan, C. Reardon, U. Woehlbier, R. H. Brown, M. Sena-Esteves, C. Gonzalez-Billault, B. Morales, L. Plate, L. W. Ruddock, M. L. Concha, C. Hetz, A. Tolun, EMBO J 2022, 41 (2), e105531, 10.15252/embj.2020105531.

[62] B. K. Sato, D. Schulz, P. H. Do, R. Y. Hampton, Mol Cell 2009, 34 (2), 212, 10.1016/j.molcel.2009.03.010.

[63] C. Huang, C. Lu, T. A. Springer, Proc Natl Acad Sci U S A 1997, 94 (7), 3156, 10.1073/pnas.94.7.3156.

[64] C. Huang, T. A. Springer, Proc Natl Acad Sci U S A 1997, 94 (7), 3162, 10.1073/pnas.94.7.3162.

[65] K. Akuta, K. Kiyomizu, H. Kashiwagi, S. Kunishima, N. Nishiura, F. Banno, K. Kokame, H. Kato, Y. Kanakura, T. Miyata, Y. Tomiyama, J Thromb Haemost 2020, 18 (2), 497, 10.1111/jth.14678.

[66] Y. Imai, E. J. Park, D. Peer, A. Peixoto, G. Cheng, U. H. von Andrian, C. V. Carman, M. Shimaoka, Blood 2008, 112 (13), 5007, 10.1182/blood-2008-03-144543.

[67] M. Semmrich, A. Smith, C. Feterowski, S. Beer, B. Engelhardt, D. H. Busch, B. Bartsch, M. Laschinger, N. Hogg, K. Pfeffer, B. Holzmann, J Exp Med 2005, 201 (12), 1987, 10.1084/jem.20041850.

[68] M. J. Feige, L. M. Hendershot, Mol Cell 2013, 51 (3), 297, 10.1016/j.molcel.2013.07.013.

[69] M. E. Call, K. W. Wucherpfennig, Mol Immunol 2004, 40 (18), 1295, 10.1016/j.molimm.2003.11.017.

[70] N. Soetandyo, Q. Wang, Y. Ye, L. Li, J Cell Sci 2010, 123 (Pt 7), 1031, 10.1242/jcs.059758.

[71] A. Blees, K. Reichel, S. Trowitzsch, O. Fisette, C. Bock, R. Abele, G. Hummer, L. V. Schafer, R. Tampe, Sci Rep 2015, 5, 17341, 10.1038/srep17341.

[72] J. M. Younger, L. Chen, H. Y. Ren, M. F. Rosser, E. L. Turnbull, C. Y. Fan, C. Patterson, D. M. Cyr, Cell 2006, 126 (3), 571, 10.1016/j.cell.2006.06.041.

[73] M. J. Shurtleff, D. N. Itzhak, J. A. Hussmann, N. T. Schirle Oakdale, E. A. Costa, M. Jonikas, J. Weibezahn, K. D. Popova, C. H. Jan, P. Sinitcyn, S. S. Vembar, H. Hernandez, J. Cox, A. L. Burlingame, J. L. Brodsky, A. Frost, G. H. Borner, J. S. Weissman, Elife 2018, 7, 10.7554/eLife.37018.

[74] S. Tian, Q. Wu, B. Zhou, M. Y. Choi, B. Ding, W. Yang, M. Dong, Cell Rep 2019, 28 (10), 2517, 10.1016/j.celrep.2019.08.006.

[75] S. Reitberger, P. Haimerl, I. Aschenbrenner, J. Esser-von Bieren, M. J. Feige, J Biol Chem 2017, 292 (19), 8073, 10.1074/jbc.M117.782284.

[76] S. Rival-Gervier, M. Y. Lo, S. Khattak, P. Pasceri, M. C. Lorincz, J. Ellis, Mol Ther 2013, 21 (8), 1536, 10.1038/mt.2013.131.

[77] F. A. Ran, P. D. Hsu, J. Wright, V. Agarwala, D. A. Scott, F. Zhang, Nat Protoc 2013, 8 (11), 2281, 10.1038/nprot.2013.143.

[78] R. Fassler, M. Meyer, Genes Dev 1995, 9 (15), 1896, 10.1101/gad.9.15.1896.

[79] U. A. Betz, C. A. Vosshenrich, K. Rajewsky, W. Muller, Curr Biol 1996, 6 (10), 1307, 10.1016/s0960-9822(02)70717-3.

[80] T. Okamoto, R. B. Schwab, P. E. Scherer, M. P. Lisanti, Curr Protoc Cell Biol 2001, Chapter 5, Unit 5 4, 10.1002/0471143030.cb0101s05.

[81] S. S. Guo, T. Y. K. Au, S. Wynn, A. Aszodi, D. Chan, R. Fassler, K. S. E. Cheah, Development 2020, 147 (22), 10.1242/dev.192724.

[82] S. B. Azimifar, R. T. Bottcher, S. Zanivan, C. Grashoff, M. Kruger, K. R. Legate, M. Mann, R. Fassler, J Cell Sci 2012, 125 (Pt 2), 435, 10.1242/jcs.091652.

[83] E. Y. Huang, M. To, E. Tran, L. T. A. Dionisio, H. J. Cho, K. L. M. Baney, C. I. Pataki, J. A. Olzmann, Mol Biol Cell 2018, 29 (9), 1021, 10.1091/mbc.E17-08-0514.

