## Supplementary data for "Transmembrane-domain-driven heterodimer assembly licenses integrin head-domain folding in the endoplasmic reticulum"

**This Supporting file contains:**

1. Supplementary figures: S1-S5
2. Key Resources Table: Antibodies, Plasmids, Cell lines, Chemicals and *etc.*

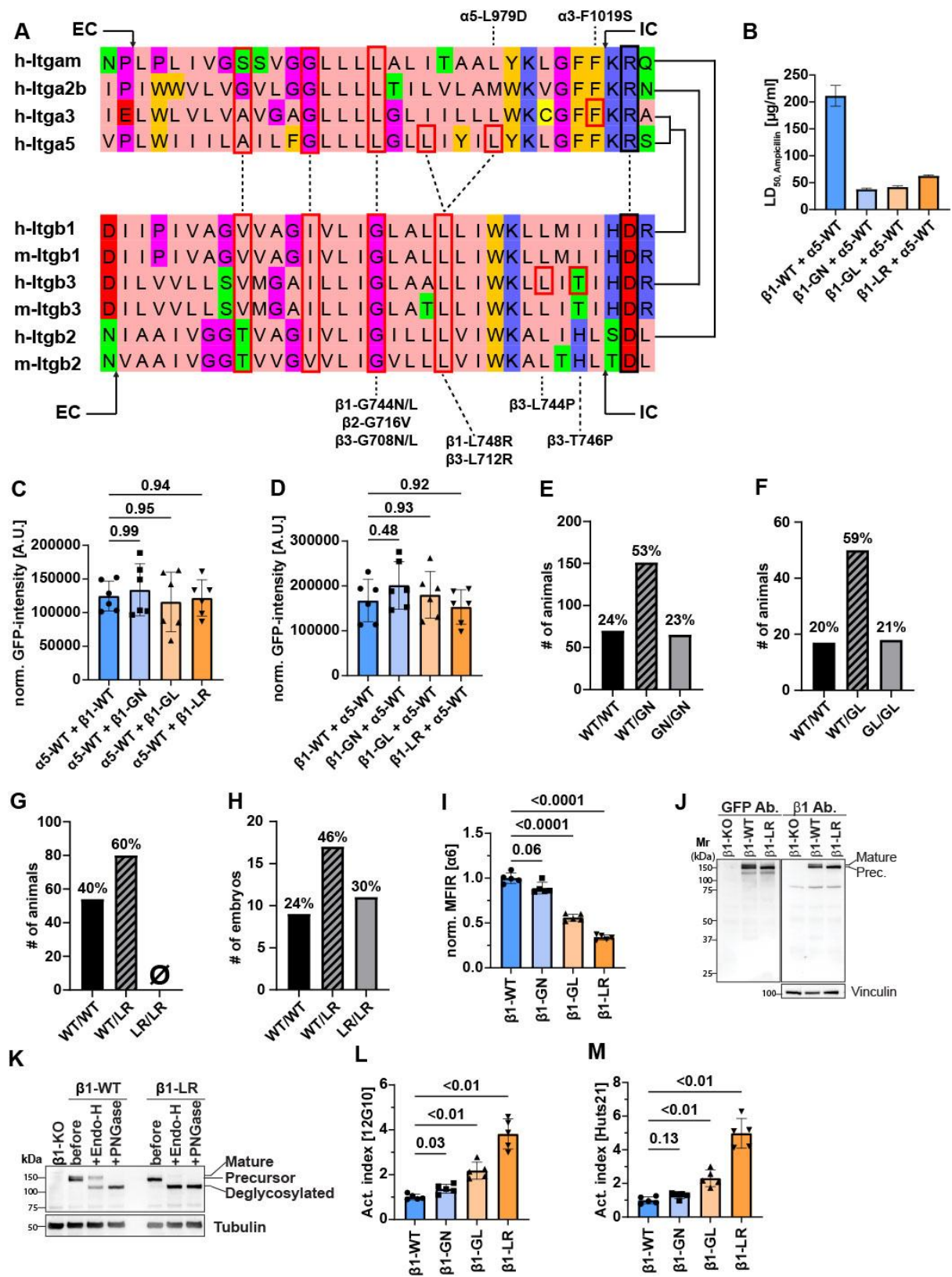

27

28

29

**Figure S1: Functional consequences of $\beta$ 1 TMD domain mutations.**

(A) Sequence alignments showing transmembrane domain residues and adjacent amino acids (EC = extracellular, IC = intracellular) of human and mouse α- and β-subunits that were investigated in the present study. Dashed lines indicate interacting amino acids on opposing α- and β-subunits based on PDB entry 2K9J. Boxed residues indicate conserved and mutated positions. Solid black lines on the right indicate interacting α-/β-subunit TMD pairs. Dashed lines above and below alignments mark mutated residues and their corresponding amino acid in each subunit. Amino acid numbering varies with signal peptide inclusion and the species (human or mouse) referenced. Alignments were generated using ClustalO (standard settings) and visualized with JalView.

(B) Bla<sup>TM</sup>-based measurements of heterotypic TMD interactions between wild-type or mutant β1 subunits fused to the C-terminal Bla and the α5 subunit fused to the N-terminal Bla in *E. coli*. Bacterial survival is reported as LD<sub>50</sub> values obtained from global fits of pooled OD<sub>600</sub> measurements from an ampicillin titration series (n=6 biological replicates).

(C,D) GFP-expression of the indicated BLA-TMD constructs in *E. coli* confirms similar expression levels of wild-type and mutant constructs. The order of subunits in the x-axis reflects the fusion of the TMD with C-BLa or N-BLa, respectively. Statistical analysis was performed using one-way ANOVA with Dunnett post-hoc test. Values represent mean±SD.

(E-H) Genotype distribution of β1 mutant mouse strains at P21 (E-G) and of β1-LR at E7.5 (H).

(I) α6 expression level analyzed by flow cytometry in cells expressing the indicated β1-subunits (n=5). Statistical analysis was performed using Brown-Forsythe and Welch ANOVA with Dunnett T3's post-hoc test. Data are presented as mean±SD.

(J) Representative Western blot of lysates from cells expressing the indicated β1-subunits. Following Western blot for GFP, the membrane was stripped and probed for β1 using the mAb Clone 18.

(K) Cell lysates from cells expressing the indicated β1-subunits were subjected to Endo-H or PNGase F digestion followed by Western blot analysis. Lysate prior to enzyme digestion served as the input control.

(L,M) Flow cytometry of cells expressing the indicated β1-subunits (n=5). The activation index was calculated as the ratio of 12G10 or Huts21 signals to total β1 signal. Statistical analysis was performed using Brown-Forsythe and Welch ANOVA with Dunnett T3's post-hoc test. Data are presented as mean±SD.

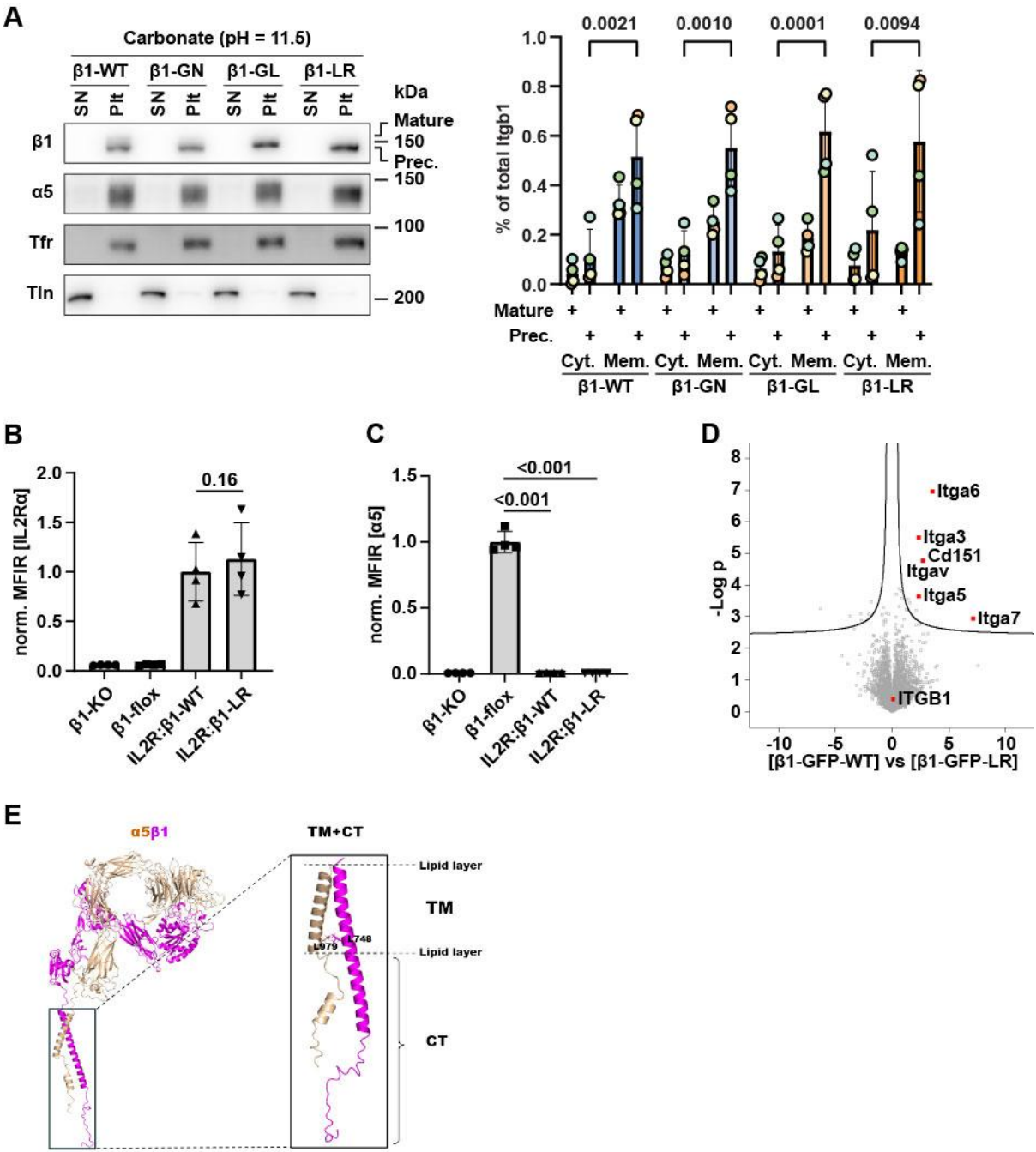

70

71

72

73

74

75

76

**Figure S2: Mutations in the $\beta 1$ TMD impair the heterodimerization with $\alpha$ -subunits.**

(A) Western blots of carbonate-extracted membrane and cytoplasmic fractions from cells expressing the indicated $\beta 1$-subunits, stained for $\beta 1$- and $\alpha 5$-integrin. Transferrin receptor (Tfr) and talin (Tln) staining served as membrane and cytoplasmic control, respectively (n=4). SN: supernatant; Plt: pellet. Densitometric quantification of precursor and mature $\beta 1$-integrin in the cytoplasmic and total membrane fraction is shown on the right as a percentage of total $\beta 1$ integrin from n=4 independent experiments.

(B,C) Flow cytometry of parental $\beta 1^{flox/flox}$ and *Cre*-deleted $\beta 1$-KO fibroblasts and $\beta 1$-KO cells reconstituted with IL2R:$\beta 1$-WT or IL2R:$\beta 1$-LR, stained for the IL2Rα ectodomain and $\alpha 5$-subunit (n=4). Statistical analysis was performed using repeated measures ANOVA with Tukey's post-hoc test. Data are presented as mean±SD.

(D) Volcano plot showing the difference between proteins co-precipitated by GFP-Trap beads from $\beta 1$-KO cells expressing $\beta 1$-WT or $\beta 1$-LR (n=4). Statistical analysis was performed using two-sided permuted t-test with 250 randomizations.

(E) Structural model of full-length $\alpha 5\beta 1$ integrin (left) predicted by AlphaFold2-multimer program and visualized in PyMOL. The side chains of $\alpha 5$-L979 and $\beta 1$-L748 in the TMD (right) are indicated.

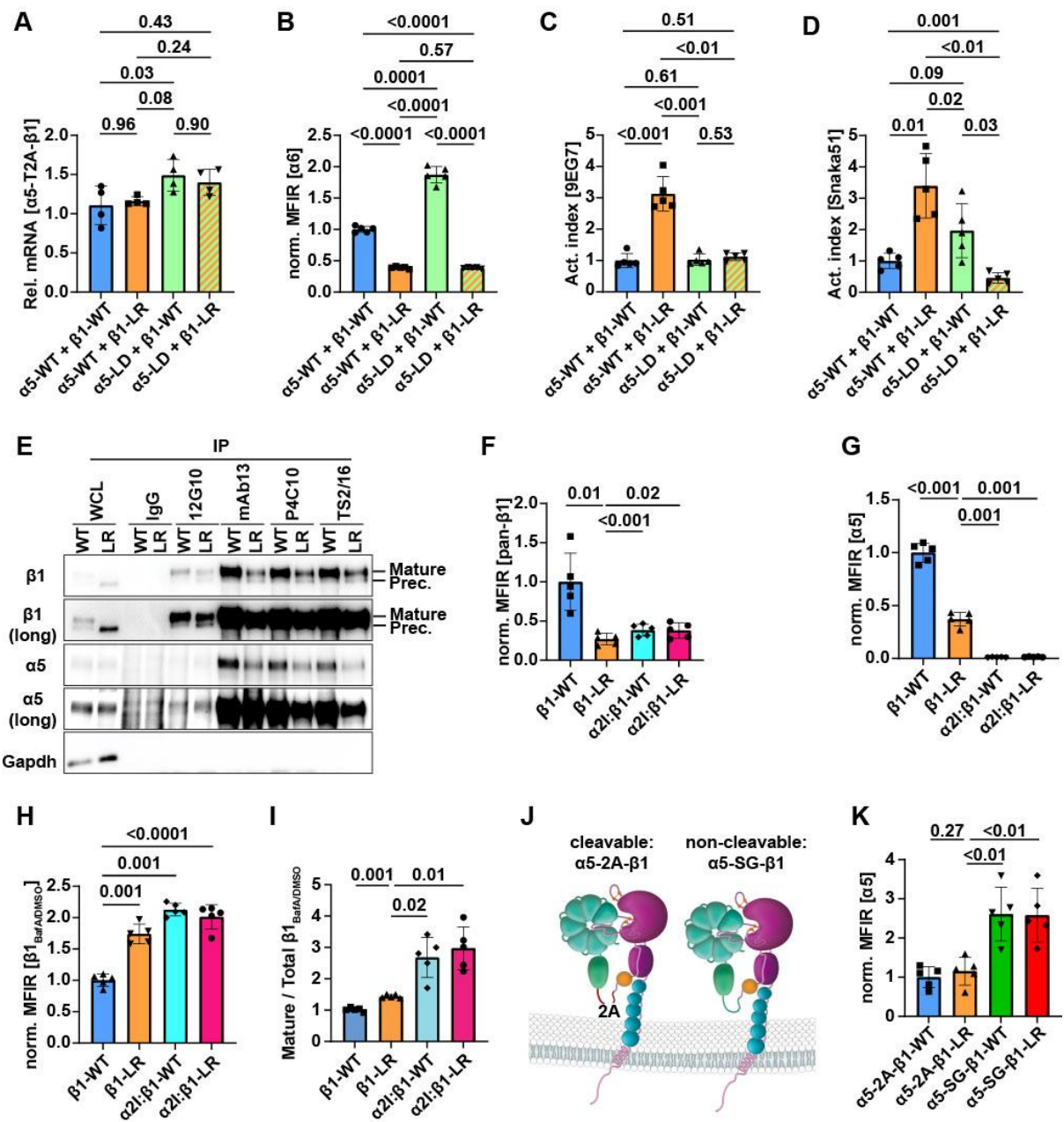

**Figure S3: The $\beta$ 1-domain of $\beta$ 1-LR causes the ER retention.**

(A) Quantitative PCR analysis of mRNA levels of the indicated α5- and β1-subunits, expressed from a single open-reading frame with a 2A self-cleaving peptide, normalized to actin in DKO cells.

(B-D) Flow cytometry-based measurements of the MFIR of α6 (B) and the activation index of the α5β1 integrin in cells expressing the indicated α5- and β1 subunits (n=5).

(C-D) The MFIR of α6 is normalized to α5β1-WT-expressing cells. Integrin activation indices were determined by calculating the ratio of the β1-activation-specific 9EG7 signal to total β1 (clone: Ha2/5, C) and the ratio of the α5-activation-specific SNAKA51 to total α5 (clone: IIA1, D). Statistical analysis for panels A-D was performed using repeated-measures ANOVA with Tukey's post-hoc test. Data are presented as mean±SD.

(E) Western blot of anti-β1 immunoprecipitates from lysates of cells expressing human β1-WT or β1-LR. Immunoprecipitations were performed with control IgG and the indicated mAbs. As a complementary experiment to Figure 3F, β1-WT and β1-LR samples were loaded side by side to facilitate direct comparison. Gapdh served as the loading control.

(F,G) Flow cytometry-based measurements of MFIR of total β1 (E) and α5 (F) in cells expressing β1-WT or β1-LR or α2I:β1-WT or α2I:β1-LR (n=5). MFIR values are normalized to β1-WT cells. Statistical analysis was performed using repeated measures ANOVA with Dunnett's post-hoc test. Data are presented as mean±SD.

(H,I) Quantification of the effect of Bafilomycin A treatment versus DMSO control on MFIR of β1 surface expression (K) and mature/total β1 integrin ratio (H) from Western blots. Data are normalized to β1-WT (n=5 for both panels). Statistical analysis was performed using repeated measures ANOVA with Dunnett's post-hoc test. Data are presented as mean±SD.

(J) Schematic representation of head and thigh domains of α5 linked to the N-terminus of β1-WT or β1-LR via either a 2A self-cleaving peptide (α5-2A-β1) or a non-cleavable SG-linker peptide (α5-SG-β1).

(J) Flow cytometry-based measurements of MFIR of α5 in cells expressing α5-2A-β1-WT or α5-2A-β1-LR, and α5-SG-β1-WT, or α5-SG-β1-LR. MFIR values are normalized to α5-SG-β1 cells.

Statistical analysis was performed using repeated measures one-way ANOVA with Dunnett's post-hoc test. Data are presented as mean±SD.

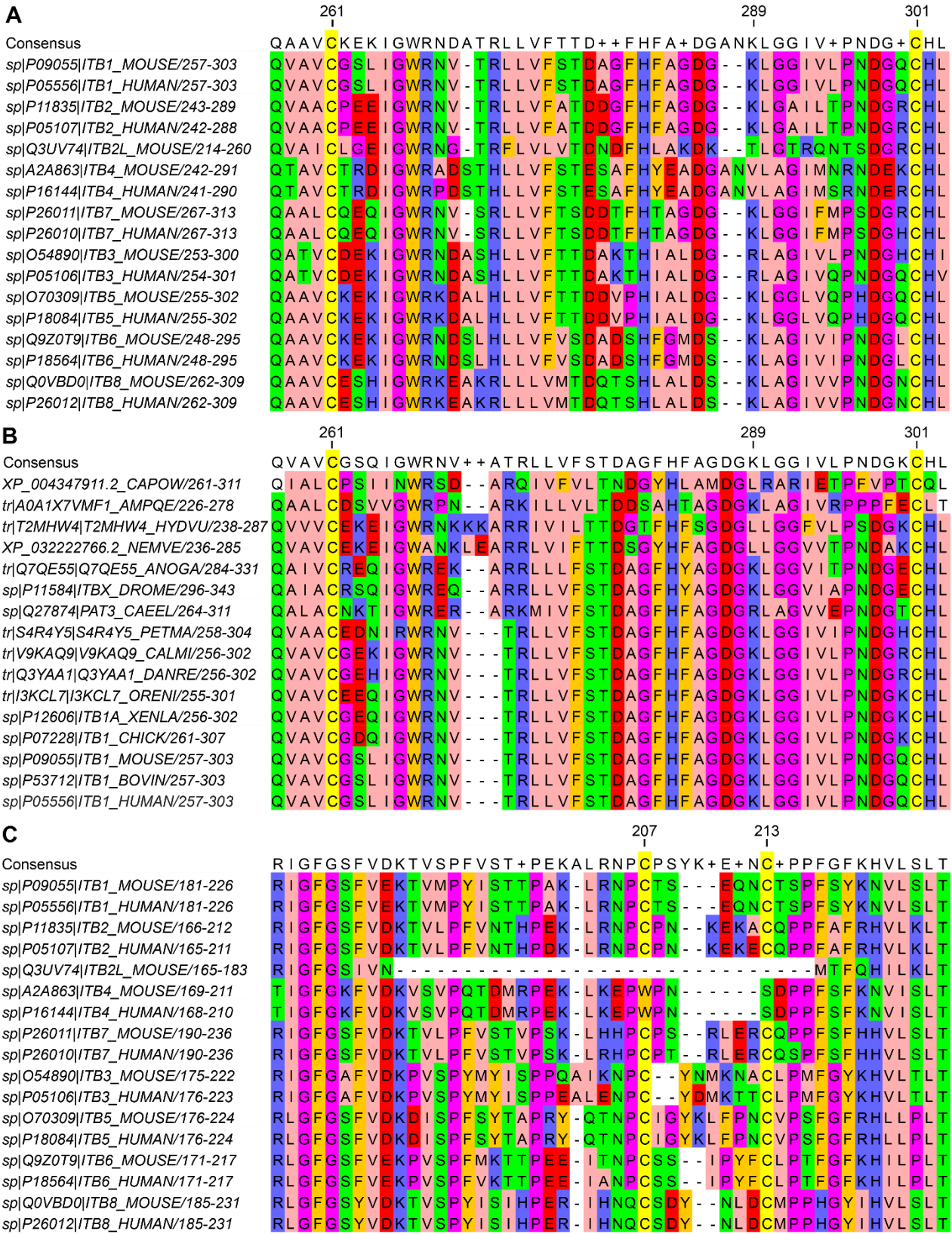

**Figure S4: C261 and C301 are conserved throughout metazoans.**

(A,B) Sequence alignments showing residues C261 and C301 in β1-subunit classes of human and mouse (A) and β1-subunits of metazoans and pre-metazoan organisms (B).

(C) Sequences alignments showing residues C207 and C213 in β1-subunit classes of human and mouse and human, as well as in the β-integrin-like Pactolus expressed in mouse granulocytes.

Alignments were performed using ClustalO (standard settings) and visualized with JalView.

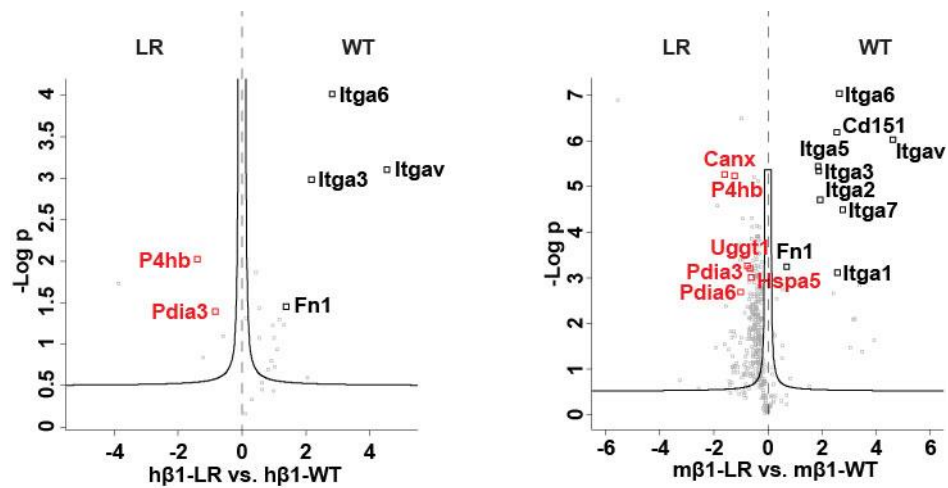

**Figure S5: Interactome analysis on β1-WT and β1-LR expressing cell lysates.**

Volcano plot of proteins co-precipitated in the cysteine oxidation mapping experiments with human β1-WT or -LR, following immunoprecipitation with the anti-human β1 mAb JB1A (A), or mouse β1-WT or -LR, following immunoprecipitation with a homemade polyclonal antibody against mouse β1 (B). Data are from 4 biological replicates. Statistical analysis was performed using two-sided permuted *t*-test with 250 randomizations. Protein disulfide isomerases enriched in β1-LR cells are highlighted in red.

**Key resources table**

| REAGENT or RESOURCE | SOURCE | IDENTIFIER |
| --- | --- | --- |
| **Antibodies** |  |  |
| β1-integrin PE (Ha2/5, 1:200) | BD Pharmingen | 562154 |
| β1-integrin PE (TS2/16, 1:200) | Biolegend | 303003 |
| β1-integrin (9EG7, FC: 1:100, IF: 1:50) | BD Pharmingen | 550531 |
| β1-integrin (12G10, 1:500) | Chemicon | MAB2247 |
| β1-integrin (Huts-21, 1:500) | BD Pharmingen | 556047 |
| mouse α5-integrin A647 (5H10-27, 1:200) | BD Pharmingen | 564312 |
| human α5-integrin A647 (IIA1, 1:200) | BD Pharmingen | 563578 |
| α6-integrin PE (GoH3, 1:100) | BD Pharmingen | 561894 |
| goat-anti-rat A647 (1:500) | Invitrogen | A21247 |
| donkey-anti-mouse A647 (1:500) | Invitrogen | A31571 |
| αM-integrin PE (M1/70, 1:50) | eBioscience | 12-0112 |
| α3-integrin (ASC-1, 1:500) | BioLegend | 343802 |
| α5-integrin (Snaka51, 1:500) | Merck | MABT201 |
| human IL2Rα/CD25 (1:500) | ThermoFisher Scientific | MA5-23714 |
| human IL2Rα/CD25 (1:500) | ThermoFisher Scientific | MA5-12680 |
| human CD71/Tfr (1:500) | BD Biosciences | 566724 |
| β3-integrin biotin (10E5, 1:500) | Novus Biologicals | NBP2-81080B |
| Streptavidin APC (1:500) | Biolegend | 405207 |
| β-actin (1:2000 for WB) | Sigma | A-2066 |
| human β1-integrin (Clone 18, 1:1000 for WB) | BD Pharmingen | 610468 |
| β1-integrin tail (1:1000 for WB) | Proteintech | 26918-1-AP |
| α5-integrin (1:1000 for WB) | Cell Signaling | #4705 |
| mouse talin-1/2 (1:1000 for WB) | Sigma | T3287 |
| mouse transferrin receptor (1:1000) | ThermoFisher Scientific | 13-6800 |
| mouse tubulin (1:1000) | DSHB | Clone: E7 |
| GFP (1:2000) | Sigma/Roche | 11814460001 |
| mouse GAPDH (1:1000) | Merck | CB1001 |
| mouse Erlec1 (1:1000) | Novus Biologicals | NBP1-86802 |
| mouse Os9 (1:1000) | Abcam | ab109510 |
| mouse Sel1l (1:1000) | Abcam | ab78298 |
| mouse Hrd1 (1:1000) | Proteintech | 13473-1-AP |
| human β1-integrin (IF: 4B7R, 1:1000) | Novus Biologicals | MAB1778 |
| mouse calnexin (IF: 1:2000) | Abcam | ab22595 |
| mouse fibronectin (IF: 1:500) | Merck | AB2033 |
| human IL2Rα/CD25 (IF: 1:1000, FC: 1:500) | ThermoFisher Scientific | MA5-23714 |
| human IL2Rα/CD25 (1:1000, FC: 1:500) | ThermoFisher Scientific | MA5-12680 |
| human α5 (1:500) | BD Pharmingen | 555651 |
| α5-tail specific antibody (IF: 1:1000) | Abcam | EPR7854 |
| mouse β1-integrin (1:10000). | Homemade, produced in rabbit | (Azimifar *et al.*, 2012) |
| 9EG7 (IP) | BD Pharmingen | 553715 |
| 8E3 (IP) | Merck | MABT199 |
| N29 (IP) | Merck | MAB2252 |
| Huts-4 (IP) | Merck | MAB2079Z |
| Huts-21 (IP) | BD Pharmingen | 556047 |
| 12G10 (IP) | Merck | MAB2247 |
| mAB13 (IP) | BD Pharmingen | 552828 |
| P4C10 (IP) | Merck | MAB1987Z |
| TS2/16 (IP) | Biolegend | 303035 |
| JB1A (IP) | Merck | MAB1965 |
| **Bacterial and virus strains** |  |  |
| *E. coli* K12 JM83 | (Schanzenbach *et al.*, 2017) |  |
| Retrovirus | In-house, see (Bottcher *et al.*, 2012) |  |
| Adeno-Cre-GFP | In-house, see (Bottcher *et al.*, 2012) |  |
| *E. coli* OmniMAX™ | ThermoFisher | C854003 |
| **Deposited data** |  |  |
| GFP or β1-GFP-WT/LR bound proteins | This study | <u>**INSERT XPrize**</u> |
| JB1A immunoprecipitated β1-WT/LR peptides | This study | <u>**INSERT XPrize**</u> |
| **Experimental models: Cell lines** |  |  |
| β1-flox, adult mouse kidney fibroblasts | (Bottcher *et al.*, 2012) |  |
| β1-KO, adult mouse kidney fibroblasts | (Bottcher *et al.*, 2012) |  |
| α5β1-dKO54 | This study |  |
| α5β1-dKO92 | This study |  |
| K562 | DSMZ | ACC-10 |
| CHO-K1 | (De Melker *et al.*, 1997) |  |
| **Experimental models: Transgenic animals/mice** |  |  |
| ITGB1 L748R (M11r) | This study |  |
| ITGB1 G744L (M10b) | This study |  |
| ITGB1 G744N (M10r) | This study |  |
| **Oligonucleotides** |  |  |
| α5-sgRNA #1 cloning oligo forward | caccgCAAGATGAGTT<br>CAGCCGATT |  |
| α5-sgRNA #1 cloning oligo forward | aaacAATCGGCTGAA<br>CTCATCTTGc |  |
| α5-sgRNA #2 cloning oligo forward | caccgCCAGATCTCG<br>TCCTCCTCGA |  |
| α5-sgRNA #2 cloning oligo reverse | aaacTCGAGGAGGAC<br>GAGATCTGGc |  |
| Me15-S (G744N/L, L748R genotyping) | ACATGTGGTCAGTC<br>TTGTTAGC |  |
| Int 14-AS (G744N/L, L748R genotyping) | GTAGTAAAGATGATT<br>GGCAGCC |  |
| Mouse actin forward | AGCCATGTACGTAG<br>CCATCC |  |
| Mouse actin reverse | CTCTCAGCTGTGGT<br>GGTGAA |  |
| Mouse GAPDH forward | TCCTGCACCACCAA<br>CTGCTTAGC |  |
| Mouse GAPDH reverse | TGGATGCAGGGATG<br>ATGTTCTGG |  |
| α5-T2A-β1 forward #1 | AGGGCAAACGTGTG<br>AGATGT |  |
| α5-T2A-β1 reverse #1 | GGGACACAGGATCA<br>GGTTGG |  |
| α5-T2A-β1 forward #2 | GAGTTTGGCCGATT<br>TGGCAG |  |
| α5-T2A-β1 reverse #2 | GGGCCCCCAGGAAA<br>TACAAA |  |
| β2-IRES2-αM forward #1 | CAGTGCCACGTTGT<br>GAGTTG |  |
| β2-IRES2-αM reverse #1 | TAGTCGCACTGGTA<br>GAGGCT |  |
| β2-IRES2-αM forward #2 | GACGGTCATGAACC<br>CCAAGT |  |
| β2-IRES2-αM reverse #2 | CGAGAGGGGAAAGA<br>CCCCTA |  |
| Chinese hamster actin forward #1 | TGACGATATCGCTG<br>CGCTC |  |
| Chinese hamster actin reverse #1 | CAGTTGGTGACAAT<br>GCCGTG |  |
| Chinese hamster actin forward #2 | CCCAAGGCCAACCG<br>TGAAAAG |  |
| Chinese hamster actin reverse #2 | TAAAGCTGTAGCCA<br>CGCTCG |  |
| **Recombinant DNA** |  |  |
| pLZRS-β1-ecto-eGFP-WT | This study | N/A |
| pLZRS-β1-ecto-eGFP-G744N | This study | N/A |
| pLZRS-β1-ecto-eGFP-G744L | This study | N/A |
| pLZRS-β1-ecto-eGFP-L748R | This study | N/A |
| pLZRS-m-Iltgb1-WT | (Bottcher *et al.*, 2012) |  |
| pRetroQ-N1-IL2Rα-β1-WT | This study | N/A |
| pRetroQ-N1-IL2Rα-β1-LR | This study | N/A |
| pRetroQ-N1-h-Iltga5-eGFP | This study | N/A |
| pRetroQ-N1-h-Iltga5-Δhead-eGFP | This study | N/A |
| HSC1-hα5-WT-T2A-hβ1-WT | This study | N/A |
| HSC1-hα5-L979D-T2A-hβ1-WT | This study | N/A |
| HSC1-hα5-WT-T2A-hβ1-L748R | This study | N/A |
| HSC1-hα5-L979D-T2A-hβ1-L748R | This study | N/A |
| pX459 | (Ran *et al.*, 2013) |  |
| N-Bla-hβ1-WT | This study | N/A |
| N-Bla-hβ1-G744N | This study | N/A |
| N-Bla-hβ1-G744L | This study | N/A |
| N-Bla-hβ1-L748R | This study | N/A |
| C-Bla-α5-WT | This study | N/A |
| C-Bla-hβ1-WT | This study | N/A |
| C-Bla-hβ1-G744N | This study | N/A |
| C-Bla-hβ1-G744L | This study | N/A |
| C-Bla-hβ1-L748R | This study | N/A |
| N-Bla-α5-WT | This study | N/A |
| N-Bla-GpA-WT | (Schanzenbach *et al.*, 2017) |  |
| C-Bla-GpA-WT | (Schanzenbach *et al.*, 2017) |  |
| N-Bla-GpA-G83I | (Schanzenbach *et al.*, 2017) |  |
| C-Bla-GpA-G83I | (Schanzenbach *et al.*, 2017) |  |
| N-Bla-hβ3-WT | This study | N/A |
| N-Bla-hβ3-L744P | This study | N/A |
| N-Bla-hβ3-T746P | This study | N/A |
| C-Bla-αIIb-WT | This study | N/A |
| C-Bla-hβ3-WT | This study | N/A |
| C-Bla-hβ3-L744P | This study | N/A |
| C-Bla-hβ3-T746P | This study | N/A |
| N-Bla-αIIb-WT | This study | N/A |
| C-Bla-α3-WT | This study | N/A |
| C-Bla-α3-GFSKR | This study | N/A |
| N-Bla-α3-WT | This study | N/A |
| N-Bla-α3-GFSKR | This study | N/A |
| C-Bla-αM-WT | This study | N/A |
| N-Bla-β2-WT-Nterm | This study | N/A |
| N-Bla-β2-G716V-Nterm | This study | N/A |
| N-Bla-β2-WT-Cterm | This study | N/A |
| N-Bla-β2-G716V-Cterm | This study | N/A |
| HSC1-hβ1-WT-i-P | This study | N/A |
| HSC1-hβ1-L748R-i-P | This study | N/A |
| HSC1-α2-αI-β1-WT-i-P | This study | N/A |
| HSC1-α2-αI-β1-L748R-i-P | This study | N/A |
| HSC1-mTm-Tfr-α2-αI-i-P | This study | N/A |
| HSC1-mTm-Tfr-β1-βI-i-P | This study | N/A |
| HSC1-mTm-Tfr-i-P | This study | N/A |
| HSC1-hβ2-WT-i-haM-WT | This study | N/A |
| HSC1-hβ2-G716V-i-haM-WT | This study | N/A |
| HSC1-α3-WT-T2A-GFP-i-P | This study | N/A |
| HSC1-α3-GFSKR-T2A-GFP-i-P | This study | N/A |
| HSC1-hβ1-CCSS-WT-i-P | This study | N/A |
| HSC1-hβ1-CCSS-L748R-i-P | This study | N/A |
| HSC1-αIIb-WT-T2A-GFP-P2A-β3-WT | This study | N/A |
| HSC1-αIIb-WT-T2A-GFP-P2A-β3-L744P | This study | N/A |
| HSC1-αIIb-WT-T2A-GFP-P2A-β3-T746P | This study | N/A |
| HSC1-ha5-head-T2A-hβ1-WT-i-P | This study | N/A |
| HSC1-ha5-head-T2A-hβ1-L748R-i-P | This study | N/A |
| HSC1-ha5-head-20SG-hβ1-WT-i-P | This study | N/A |
| HSC1-ha5-head-20SG-hβ1-L748R-i-P | This study | N/A |
| HSC1-hβ1-WT-i-ha5-WT | This study | N/A |
| HSC1-hβ1-L748R-i-ha5-WT | This study | N/A |
| HSC1-hβ1-WT-8xKR-i-ha5-4xKR | This study | N/A |
| HSC1-hβ1-L748R-8xKR-i-ha5-4xKR | This study | N/A |
| **Software and algorithms** |  |  |
| ImageJ – Fiji | (Schindelin *et al.*, 2012) | [https://imagej.net/W<br>elcome](https://imagej.net/welcome) |
| Perseus 1.6.2.3 | (Tyanova *et al.*, 2016) | https://maxquant.net/<br>perseus/ |
| Spectronaut 18.0 | Spectronaut Software | https://biognosys.com/software/spectronaut/ |
| MaxQuant (version differs, see methods) | (Cox and Mann, 2008) | https://www.maxquant.org/ |
| Origin (Pro), 2023 | OriginLab Corporation | https://www.originlab.com/ |
| Biorender | Biorender online software | https://www.biorender.com/ |
| GraphPad Prism 9.0 / 10.0 | GraphPad Software | https://www.graphpad.com |
| Uniprot-align | The UniProt Consortium (Apweiler et al., 2004) | https://www.uniprot.org/align |
| JalView 2.11.2.6 | (Waterhouse et al., 2009) | https://www.jalview.org/download/ |
| FlowJo Software v10.6.1 | BD Biosciences | https://www.flowjo.com/solutions/flowjo/downloads |
| UCSF Chimera v.1.17.3 | (Pettersen et al., 2004) | https://www.cgl.ucsf.edu/chimera/ |
| **Chemicals, biologicals, peptides, and recombinant proteins** |  |  |
| MG132 | Sigma | 474787 |
| CB5083 | MedChemExpress | HY-12861 |
| DMSO | Serva | 2038501 |
| Bafilomycin A1 | Adipogen | BVT-0252 |
| Puromycin | Sigma | P8833 |
| Kanamycin | Sigma | 10106801001 |
| Chloramphenicol | Sigma | C0378 |
| Ampicillin | Sigma | A9518 |
| NEM | Sigma | E3876 |
| d5-NEM | Sigma | 692905 |
| Sodium deoxycholate | Sigma | 30970 |
| 2-chloroacetamide | Sigma | 148415000 |
| Tris(2-carboxyethyl)-phosphine, hydrochloride | ThermoFisher | T2556 |
| Trypsin, sequencing grade | Promega | V5111 |
| GluC, sequencing grade | Promega | V1651 |
| Trifluoroacetic acid | Merck | 80457 |
| L-(+)-Arabinose | Sigma | A3256 |
| LB-medium | Carl Roth | X964.2 |
| Isopropyl-β-d1-thiogalactopyranoside | Sigma | I5502 |
| cOmplete protease inhibitor tablets | Roche | 04693159001 |
| Triton X-100 | Roth | 6683.1 |
| CHAPS | Merck | C5070 |
| Tween-20 | Serva | 37470.1 |
| Digitonin | Merck | D141 |
| Sodium dodecylsulfate | Roth | CN30.3 |
| Bovine serum albumin | PAN Biotech | P06-1391100 |
| Gibco™ Fetal bovine serum | ThermoFisher | 10270-106 |
| Gibco™ DMEM, high glucose, GlutaMAX™ Supplement, pyruvate | ThermoFisher | 10569010 |
| Gibco™ RPMI-1640 | ThermoFisher | 11875093 |
| Gibco™ Ham's F-12 nutrient mix | ThermoFisher | 11765054 |
| Gibco™ GlutaMAX™ Supplement | ThermoFisher | 35050061 |
| Bis-Tris, powder | Sigma | B9754 |
| ε-Caproic acid, powder | Sigma | A2504 |
| 2-Mercaptoethanol | Merck | 8.05740.0250 |
| Fibronectin | Sigma | F1141 |
| Protein A/G PLUS-Agarose | SCBT | sc-2003 |
| Hoechst 33342, trihydrochloride trihydrate | ThermoFisher | H1399 |
| ibidi Mounting Medium | ibidi | 50001 |
| Ibidi Immersion Oil | ibidi | 50101 |
| μ-Slide 8 Well, ibi-treat | ibidi | 80826 |
| MS grade water | VWR | HEWL5190-6897 |
| Paraformaldehyde, powder | Merck | 104005 |
| EasyTag EXPRESS 35S Protein Labeling Mix, 7 mCi | Revvity | NEG772007MC |
| Gibco™ DMEM, high glucose, no glutamine, no methionine, no cystine | ThermoFisher | 21013024 |
| SILAC FBS (dialyzed) | Silantes | 281000900 |
| **Other** |  |  |
| ReproSil-Pur C18-AQ 1.9-micron beads | Dr. Maisch GmbH | r119.aq. |
| Easy-nLC 1200 system | ThermoFisher |  |
| Evotip Pure | Evosep | EV2018 |
| PepSep C18, 15cmx150μm, 1.5 μm | Bruker Daltonics | 1893471 |
| Evosep One HPLC | Evosep |  |
| **Critical commercial assays** |  |  |
| NEBuilder® HiFi DNA Assembly Master Mix | New England Biolabs | M5520 |
| PfuUltra II | Agilent Technologies | 600670 |
| Q5 Hot Start polymerase | New England Biolabs | M0493 |
| Pierce™ BCA Protein Assay Kits | ThermoFisher | 23225 |
| NativePAGE™ 4 to 16%, Bis-Tris, 1.0 mm, Mini Protein Gel, 15-well | ThermoFisher | BN1004BOX |
| NativePAGE™ Sample Buffer (4X) | ThermoFisher | BN2003 |
| NativePAGE™ Cathode Buffer Additive (20X) | ThermoFisher | BN2002 |
| NativePAGE™ 5% G-250 Sample Additive | ThermoFisher | BN2004 |
| NativePAGE™ Running Buffer (20X) | ThermoFisher | BN2001 |
| PVDF membrane Immobilon-P | Merck | IPVH00010 |
| Immobilon™ Western HRP substrate | Merck | WBKLS0500 |
